# Spontaneous cortical vasodynamics form a multiscale propagation architecture across the awake brain

**DOI:** 10.64898/2026.05.19.726296

**Authors:** Xiaochen Liu, Xiaoqing Zhou, David Hike, Jacob Duckworth, Nivetha Pasupathy, Yuanyuan Jiang, Soonwook Choi, Zhanyan Fu, Bruce Rosen, David Kleinfeld, Xin Yu

## Abstract

Spontaneous vasomotor rhythms and fluctuations are recognized contributors to functional MRI (fMRI) signals. Yet the spatial evolution of vasomotor contributions, which bear on the interpretation of fMRI signals in terms of functionally connected brain regions, remains unresolved. To clarify this issue, we analyze the 0.01 - 0.20 Hz space-frequency modes of 14T cerebral blood volume (CBV)-weighted fMRI across the awake mouse cortex. High-resolution CBV-fMRI identified single-vessel-aligned, frequency-specific modes of vasodynamic propagation, and ultra-fast CBV-fMRI enabled mapping of phase-gradients along tangential and radial axes. Tangential gradients extended maps of vasomotor traveling waves across the entire cortical mantle, with a median speed of 0.9 - 1.3 mm/s. Radial gradients revealed approximately three-fold slower changes in timing that support lamina-dependent regulation of perfusion. Region-specific differences in propagation speed and direction occur across cortical areas and between mice. These findings reframe vascular fluctuations as a feature of brain-wide physiology, as opposed to residual noise.

## Introduction

Functional MRI (fMRI) measures brain activity indirectly through hemodynamic signals^1–6^ whose spatiotemporal dynamics are shaped by local neurovascular coupling^3,7^, neuromodulatory regulation of vascular tone^8,9^, and intrinsic arteriole smooth muscle contractility that generates low-frequency vasomotion^10,11^. In resting-state fMRI, these spontaneous hemodynamic fluctuations have transformed the study of intrinsic brain organization by revealing reproducible functional networks, dynamic connectivity patterns, and propagating activity motifs across the brain^12–16^. However, the vascular component of these signals has a dual identity: it can obscure inference about neural connectivity, yet it also encodes organized physiological information about cerebrovascular function^2,3,8,10,17,18^. Because spontaneous vascular fluctuations arise from active processes that regulate blood flow, blood volume, vascular tone, tissue oxygenation, and fluid exchange^10,11,19–27^, a key unresolved question is whether these vasodynamics exhibit organized propagation across the awake brain.

Spontaneous vasodynamics are expected to be spatially and temporally coordinated because they arise from interacting vascular and neural mechanisms rather than from isolated local fluctuations. Vasomotion originates from intrinsic contractility of vascular smooth muscle in arterioles^28,29^, can be coordinated across neighboring vessels through endothelial K^+^-signaling^30^, and can be entrained by neuronal activity^10,31^. The associated oscillatory vascular dynamics exhibit spatial coherence and structured phase relationships across arteriolar networks^10,18,32^. At larger scales, optical imaging has shown that vasomotor oscillations propagate as long-wavelength traveling waves across the cortical surface^17^. Thus, prior work has established coordination within local vascular networks and along the cortical surface, but whether this coordination extends to three-dimensional propagation across the intact awake cortex remains unclear. Also unclear is propagation across the large number of cortical regions inaccessible to optical imaging.

Coordinated vascular dynamics may contribute to cerebrospinal fluid-interstitial fluid exchange along perivascular pathways, particularly under brain-state-dependent neuromodulation during sleep^20,24,25^. Consistent with this view, impairment of vasomotion has been shown to disrupt perivascular clearance of amyloid-β, potentially linking vasodynamics to neurodegenerative processes^21–23,33^. In parallel, neuromodulatory systems, including noradrenergic and cholinergic pathways, regulate vascular tone and coordinate vascular dynamics across the cortex, and their disruption in aging and neurodegeneration is associated with impaired vascular regulation and altered hemodynamic dynamics^26,27,34–36^. Together, these findings underscore the functional importance of coordinated vascular dynamics and suggest that brain-wide vasodynamic organization may provide a unique readout of cerebrovascular dysfunction.

Despite this emerging importance, measuring vasodynamic organization across the brain remains challenging. Conventional fMRI studies have identified disease-related alterations in low-frequency signal fluctuations and functional connectivity patterns^12,13^. More recent work has demonstrated that resting-state BOLD signals exhibit structured propagation, with phase delays forming traveling waves linked to arousal and systemic physiology^14–16^. However, the BOLD signal reflects a complex mixture of cerebral blood flow, blood volume, and oxygen metabolism, making it difficult to isolate vasodynamics from neuronal and metabolic contributions^16^. As a result, the specific spatiotemporal organization of intrinsic vasodynamics cannot be directly resolved using BOLD contrast alone. High-field cerebral blood volume (CBV)-weighted fMRI provides a more direct approach to probing vasodynamics^37–40^. In particular, single-vessel CBV fMRI^41–43^ has demonstrated that arteriolar oscillations can be resolved with vessel-level specificity and exhibit spatially structured phase relationships with a characteristic correlation length on the order of millimeters^18^. These findings establish that vasodynamic signals are accessible to MRI and carry intrinsic spatial organization. Yet whether such an organization can be mapped across the entire cortical mantle, and whether its propagation structure can be characterized in three dimensions in the awake brain, remains unknown.

In the present study, we address this gap by combining CBV-weighted fMRI with space-frequency singular value decomposition (SF-SVD)^44,45^ to map vasodynamics across the awake mouse cortex. SF-SVD enables the identification of frequency-specific coherent oscillatory modes and their associated phase structure across space^17,46^. Applying this framework to high-resolution and ultra-fast CBV-fMRI data acquired at 14 T, we demonstrate that vasodynamic activity exhibits organized spatiotemporal propagation across the cortex, with distinct frequency-dependent spatial modes and three-dimensional phase gradients. These results establish a noninvasive framework for resolving vasodynamic propagation at the systems level, opening a path toward vascular dynamic signatures of cerebrovascular dysfunction.

## Results

### SF-SVD applied to high-resolution whole-brain CBV fMRI recovers vessel-specific vasodynamic oscillations

Awake mouse fMRI was performed at 14 T using an implanted radiofrequency (RF) coil and head-fixation approach as described previously^47^. After intraperitoneal administration of MION, penetrating vessels were identified as dark lines oriented perpendicular to the cortical surface in both anatomical FLASH and raw functional EPI images (Fig. 1D–E). To assess whether CBV-based fMRI signal fluctuation presents spatially structured vasodynamic information, independent component analysis (ICA) was applied to the time series (Supp. Fig. 1A). The dominant component exhibited spatial patterns aligned with penetrating vessel locations (Fig. 1F-G) with a corresponding time course showing slow fluctuations within the vasomotion frequency range (Fig. 1H-I). These data indicate that CBV contrast provides sufficient spatial specificity to resolve vessel-associated dynamics.

**Figure 1.**
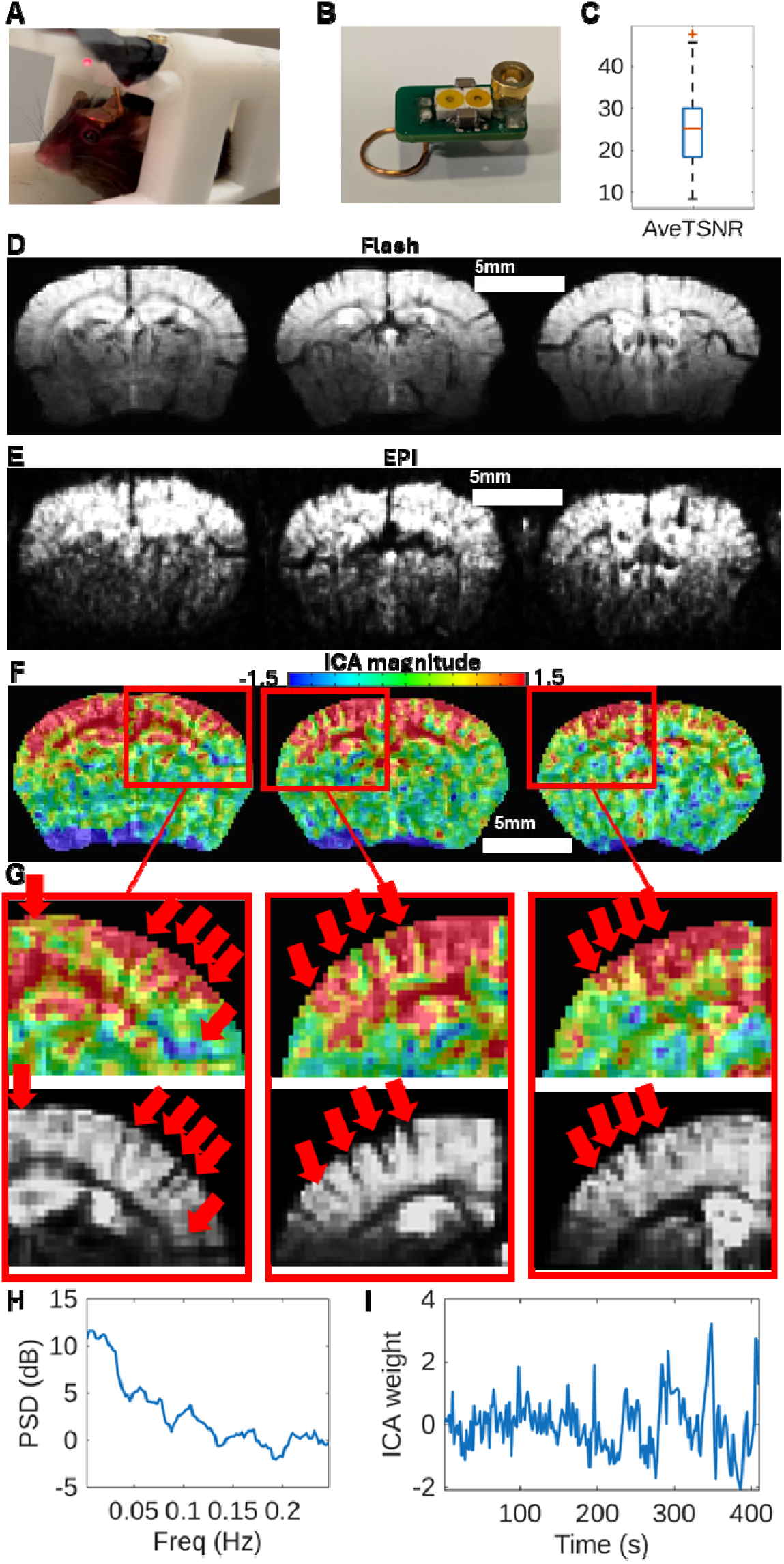
High-resolution awake mouse CBV fMRI and ICA based vessel-specific signal decomposition. (A) Photograph of the custom head-fixation holder with an awake, head-fixed mouse. (B) The surgically implanted receive coil used for 14T CBV fMRI. (C) Boxplot of the average temporal signal-to-noise ratio (TSNR) across all trials (n=8 mice, 55 trials). (D) Representative anatomical FLASH image showing cortical vascular architecture, with penetrating vessels visible as dark lines through the gray matter. (E) Corresponding CBV EPI image from the same session, with the same vessel patterns identifiable. (F) Spatial magnitude map of the dominant ICA component, showing non-random structure concentrated along the vascular architecture. (G) Zoomed-in comparison between the ICA magnitude map (top) and the anatomical FLASH image (bottom) for three representative coronal slices, demonstrating alignment of high-magnitude ICA regions with penetrating vessel positions. Red arrows indicate corresponding vessels. (H) Power spectral density of the dominant ICA component time course, showing multiple spectral peaks in the vasomotion frequency band. (I) Time course of the dominant ICA component weight over a representative 410-second trial.

Because ICA operates on broadband signals and does not separate frequency-specific contributions, we switched to space-frequency SVD (SF-SVD) to resolve the spectral structure of vasodynamic activity (Supp. Fig. 1B). SF-SVD identified distinct coherence peaks across the 0.01-0.20 Hz frequency range and extracted corresponding spatial magnitude maps for each frequency (Fig. 2). These maps revealed vessel-aligned patterns that differed across frequency ranges, indicating that vasodynamic oscillations are not uniformly distributed across the vasculature but exhibit frequency-dependent spatial selectivity (Fig. 2B-D, F-H).

**Figure 2.**
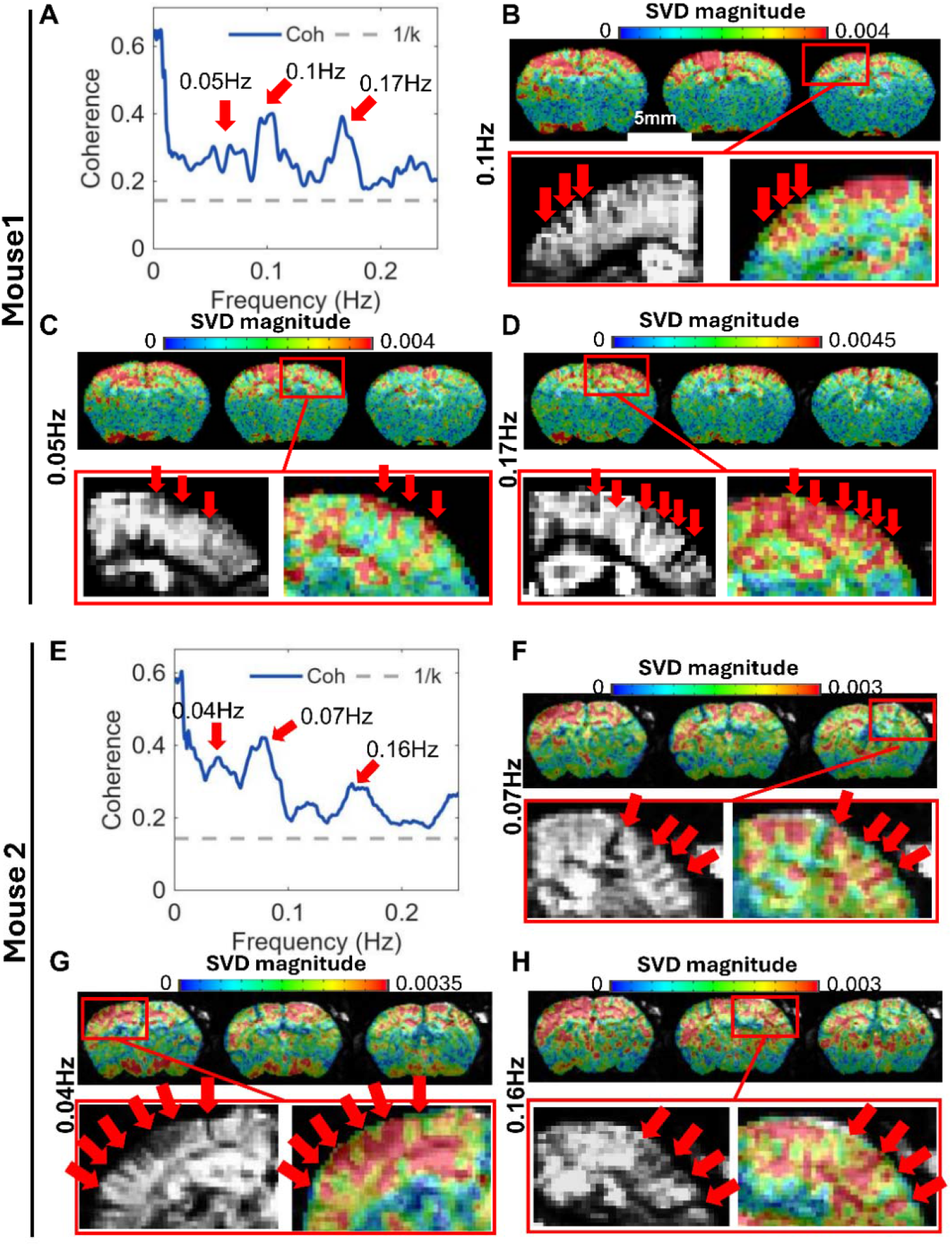
SF-SVD resolves frequency-specific vasodynamics in high-resolution CBV fMRI. (A) SF-SVD coherence spectrum from a representative trial of Mouse 1, with prominent peaks at 0.05, 0.1, and 0.17 Hz. The dashed line shows the 1/k noise floor. (B–D) SF-SVD magnitude maps of the cortical volume at 0.1 Hz (B), 0.05 Hz (C), and 0.17 Hz (D), with zoomed-in comparisons to th anatomical image showing vessel alignment at each frequency. Red arrows indicate vessel-aligned features. Scale bar, 5 mm. (E) SF-SVD coherence spectrum from a representative trial of Mouse 2, with peaks at 0.04, 0.07, and 0.16 Hz. The dashed line shows the 1/k noise floor. (F–H) Corresponding SF-SVD magnitude maps at 0.07 Hz (F), 0.04 Hz (G), and 0.16 Hz (H) for Mouse 2, with zoomed-in vessel comparisons.

We next performed population-level analysis of the spectral-spatial organization. Across all trials, SF-SVD revealed a broad distribution of center frequencies spanning 0.01-0.20 Hz without a single dominant frequency (Fig. 3A-B). Informed by the trial-based histogram distribution, frequencies were grouped into three bands (low, intermediate, and high; Fig. 3B). The resultant modes revealed systematic differences in both signal magnitude and spatial organization. In particular, cortical SF-SVD magnitude was significantly lower in the low-frequency band compared to higher-frequency bands (Fig. 3C), indicating frequency-dependent differences in the strength of coherent vascular activity. To further characterize the spatial structure underlying these differences, non-negative matrix factorization (NMF) identified three dominant components corresponding to anterior cortex, posterior cortex, and subcortical regions (Fig. 3D). Notably, cortical components were preferentially associated with intermediate and high-frequency bands, whereas the subcortical component was selectively enriched in the low-frequency band (Fig. 3E), revealing a structured coupling between frequency and anatomical organization. Together, these results demonstrate that vasodynamics exhibit a frequency-dependent spatial organization across the brain, establishing vessel-specific spectral structure as a general property of cerebrovascular dynamics in the awake state.

**Figure 3.**
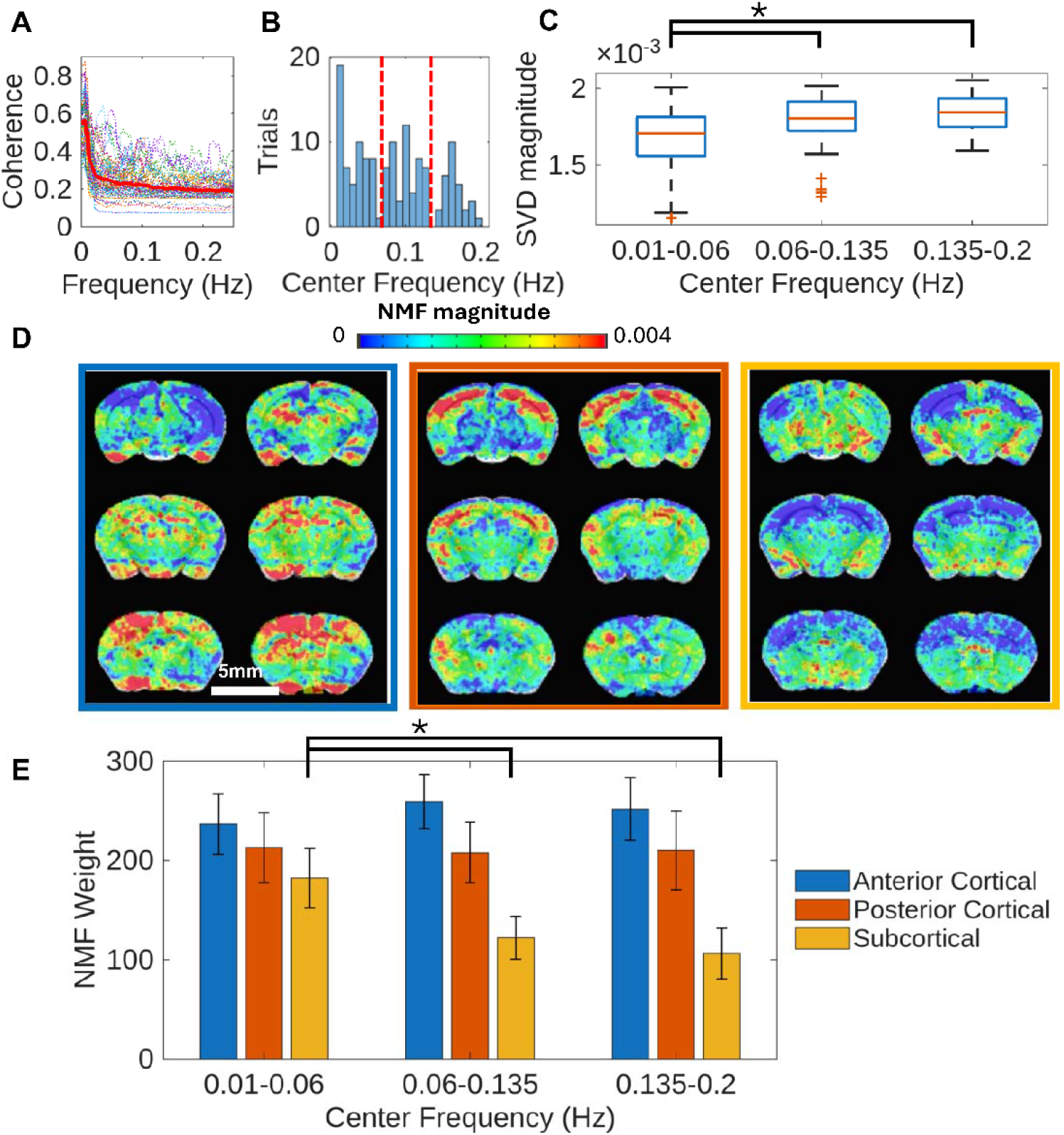
SF-SVD based group-level properties of vasodynamics. (A) SF-SVD coherence spectra from all 55 trials (colored traces) with the trial-averaged coherence (red). (B) Distribution of detected center frequencies across the 55 trials, showing broad spectral coverage from 0.01 to 0.2 Hz. Dashed red lines indicate the boundaries between frequency groups used in (C) and (E) .(C) Boxplot of the average SF-SVD magnitude within the cortical area grouped by center frequency band (0.01–0.06 Hz, 0.06–0.135 Hz, 0.135–0.2 Hz). The sub-0.06 Hz group showed significantly lower magnitude than both higher-frequency groups (*: two-sample t-test, p<0.05). (D) The three dominant NMF spatial components extracted from the magnitude maps across 55 trials, corresponding to anterior cortex (blue border), posterior cortex (orange border), and subcortical structures (yellow border). (E) NMF component weights grouped by center frequency band for the three spatial components. The subcortical component (yellow) showed significantly higher weight in the sub-0.06 Hz group compared to higher-frequency groups (*: one-way ANOVA with Tukey post-hoc, p<0.05).

### Validation of vasodynamic propagation detection with SF-SVD

As a control experiment to validate that SF-SVD accurately recovers phase information, we applied SF-SVD to a controlled bilateral forepaw stimulation paradigm in anesthetized mice. Here, the left and right somatosensory cortices were activated using a 0.025 Hz stimulation paradigm with a fixed 5 s delay (Fig. 4A). SF-SVD correctly localized activation to both cortices (Fig. 4B) and recovered a consistent interhemispheric phase offset corresponding to a median lag of 4.8 s across animals (Fig. 4C-E), in close agreement with the imposed delay. The variability likely reflects differences in vascular propagation dynamics and the distributed nature of the cerebrovascular network across hemispheres.

**Figure 4.**
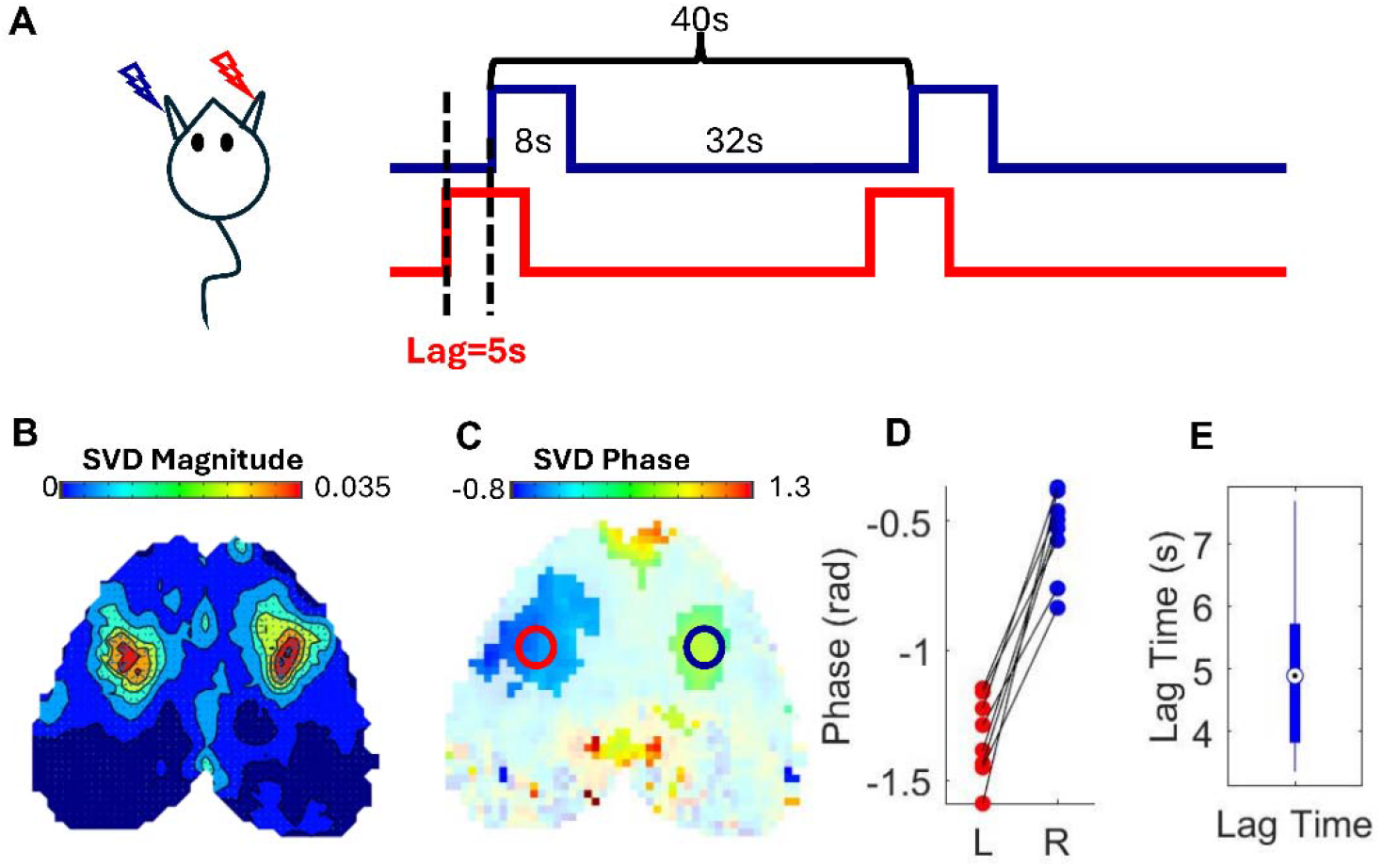
SF-SVD for inter-hemispheric phase lag detection from a controlled stimulation paradigm. (A) Schematic of the bilateral forepaw stimulation paradigm. Left and right forepaws were stimulated in alternating 8s-on-32s-off cycles, with the left side delayed by 5 seconds relative to the right, generating a 0.025 Hz CBV oscillation with a predictable inter-hemispheric timing offset. (B) Trial-averaged SF-SVD magnitude map at 0.025 Hz, with bilateral high-amplitude regions at the left and right somatosensory cortices (n=8 mice). (C) Corresponding trial-averaged SF-SVD phase map, showing the timing offset between the two somatosensory regions. Circles indicate the ROIs used for phase extraction. Here, delay increases with increasing phase. (D) Phase values extracted from left (L) and right (R) somatosensory cortex ROIs for each animal, showing a consistent phase offset with the left hemisphere (right forepaw) leading. (E) Boxplot of the estimated lag time computed as for each animal. The median lag time was 4.8 s, in close agreement with the 5-second stimulation offset.

We next examined whether SF-SVD can resolve spatial phase gradients in spontaneous vasodynamic activity using the same awake CBV fMRI dataset described above. At the temporal resolution of this acquisition (TR = 2 s), phase accumulation imposes a spatial resolution limit of several millimeters for reliable phase estimation (Supp. Fig. 2A), precluding continuous voxel-wise mapping, particularly along the cortical depth. Instead, phase gradients were estimated within ∼4 mm cortical regions aligned with vessel structures identified in the SF-SVD magnitude maps (Supp. Fig. 2B-C). These analyses yielded consistent cortical phase gradients on the order of ∼0.4-0.5 rad/mm and corresponding propagation speeds of 2.3 ± 0.02 mm/s across trials (Supp. Fig. 2D), in agreement with prior optical measurements^17^. To assess whether these propagation features generalize beyond CBV fMRI, we applied SF-SVD to functional ultrasound (fUS) data acquired under light sedation. Similar vasodynamic signal fluctuations were observed within the same frequency range, and phase gradient estimates using the same analysis approach yielded comparable values (∼0.4 rad/mm) to those obtained with CBV fMRI (Supp. Fig. 3). Together, these results demonstrate that SF-SVD enables robust recovery of vasodynamic phase relationships and supports reliable estimation of cortical propagation gradients across modalities. These data provide a principled framework for mapping spatiotemporal vascular dynamics in vivo.

### Ultra-fast CBV fMRI enables spatially resolved 3D cortical vasodynamic propagation mapping

To overcome the temporal-sampling constraint on resolving short-range spatial phase differences imposed by the 0.5 Hz sampling rate (TR = 2 s), we increased the acquisition rate to 8 Hz (TR = 0.125 s). The shorter TR reduced the phase advance between successive samples and decreased the propagation distance per sampling interval from several millimeters to the submillimeter range (Supp. Fig. 2A), thereby enabling spatially resolved estimation of depth-wise phase gradients across the cortical mantle. This improvement brings the full cortical depth within the resolvable range and enables estimation of phase gradients along both tangential and radial axes. To facilitate this analysis, the curved cortical volume was geometrically flattened to separate tangential propagation along the cortical surface from radial, depth-dependent propagation (Supp. Fig. 4). Representative anatomical (FLASH) and functional raw EPI images from the 8 Hz acquisition are shown in Fig. 5A-B. SF-SVD identified coherent oscillatory components with center frequencies broadly distributed between 0.05 and 0.2 Hz across trials (Fig. 5C-D), consistent with the spectral structure observed in the high-resolution dataset. At each frequency, SF-SVD revealed spatially continuous phase maps that exhibited smooth gradients across the tangential cortical plane (Fig. 5E), indicating large-scale propagation patterns consistent with prior wide-field measurements of long-wavelength vasomotion traveling waves^17^.

**Figure 5.**
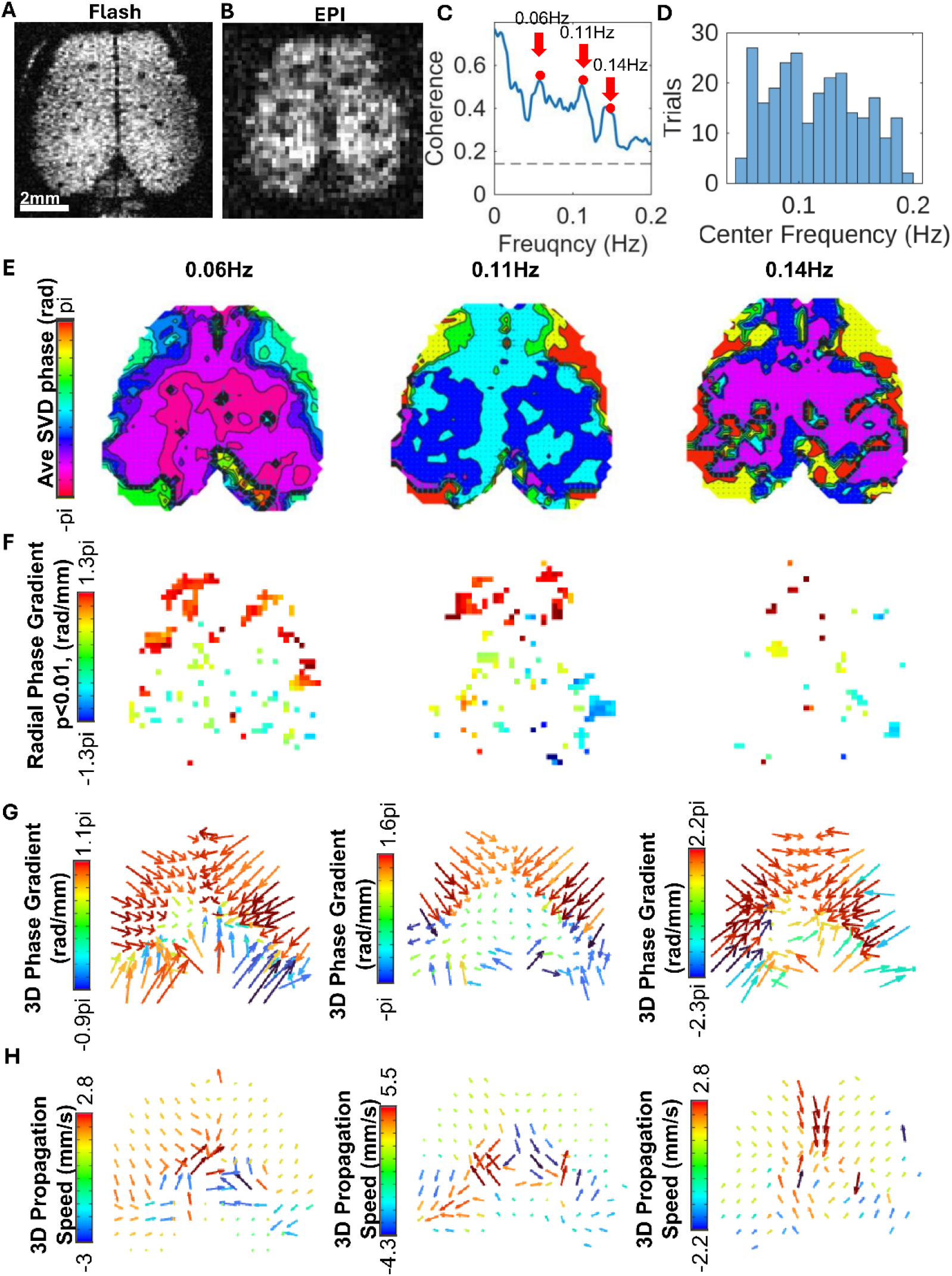
Ultra-fast CBV fMRI resolves spatially continuous 3D vasodynamic propagation across the awake mouse cortex. (A) Representative anatomical FLASH image from the awake mouse cortex. (B) Corresponding CBV EPI image. (C) SF-SVD coherence spectrum from a representative trial, with dominant peaks at 0.06, 0.11, and 0.14 Hz. (D) Distribution of detected center frequencies across all 118 trials from 18 mice. (E) Depth-averaged SF-SVD phase maps across the tangential cortical plane at 0.06 Hz, 0.11 Hz, and 0.14 Hz from the representative trial in (C), showing cortex-wide gradually varying patterns. Here, delay increases with increasing phase. (F) Radial phase gradient map, showing the depth-dependent phase gradient at each cortical position. Only voxels with a significant radial gradient (p < 0.01) are displayed. Color encodes gradient magnitude; sign encodes the radial direction. (G) 3D phase gradient vector maps at each of the three center frequencies. Vectors are downsampled to a 0.6 mm display grid. Arrow length reflects the tangential projection of the vector magnitude; color encodes total 3D gradient magnitude in rad/mm; sign encodes the radial direction. (H) 3D propagation speed vector maps at the same three frequencies, derived by converting phase gradients via *c* = 2π*f*/ g. Arrow length reflects the tangential projection of the speed magnitude; color encodes propagation speed in mm/s; sign encodes the radial direction.

Beyond the tangential plane, cortex-wide phase gradients along the local radial axis were estimated using linear regression through cortical depth. Significant gradients (p < 0.01) exhibited a sparse spatial distribution and were organized in a bidirectional manner across the cortex (Fig. 5F), with positive and negative values indicating opposite directions of depth-dependent phase progression. Representative positive and negative radial gradients showed approximately linear phase progression through cortical depth at discrete locations aligned with vessel-visible features (Supp. Fig. 5). These profiles support a laminar-specific readout of vasodynamic timing and are consistent with a localized, vessel-associated organization rather than a uniformly expressed radial pattern across the cortex. The mean phase-gradient magnitude (1.9 ± 1.2 rad/mm) was substantially larger than the typical along-vessel vasomotion gradient with mean magnitude |g| = 0.43 rad/mm^17^, suggesting that the laminar-specific radial signal may reflect contributions from multiple sources of depth-dependent vascular timing.

To integrate both tangential and radial components, full 3D phase gradient vectors were computed at each frequency (Fig. 5G). To estimate local phase variation, gradients were calculated using a sliding spatial fitting window, in which a linear model was fit using a 2 mm tangential fitting window and a radial window of at least 0.8 mm at each cortical location to derive the local phase gradient vectors. The 3D phase gradient vectors were presented in tangential view with a ±0.6 mm median downsample window. These maps revealed coherent propagation patterns across the cortex, with consistent spatial organization while exhibiting frequency-dependent scaling in magnitude.

To enable direct comparison across frequencies, phase gradients were converted to vectors whose magnitude is the propagation speeds (Fig. 5H). The resulting maps showed typical propagation speeds on the order of 1–2 mm/s across frequencies, comparable to values reported for vasomotion along pial and penetrating arterioles using two-photon microscopy^17^. However, these measurements are not directly equivalent, as the CBV fMRI signal reflects an aggregate response from a heterogeneous vascular population rather than propagation along individual arterioles. Nonetheless, these results demonstrate that vasodynamic propagation in the awake cortex exhibits a coherent three-dimensional organization, integrating vessel-scale dynamics with depth-dependent vascular timing structure.

### Group-level organization of 3D vasodynamic propagation

To characterize the group-level organization of vasodynamic propagation, we applied principal component analysis (PCA) to trial-wise 3D propagation speed maps, computed using a 2 mm sliding spatial fitting window to estimate local phase gradients. Across all trials, the three dominant components (47%, 36%, and 17% variance explained; Fig. 6A-C) revealed spatially organized propagation patterns characterized by bilateral divergence from the midline, anterior–posterior convergence, and focal divergence centered at anterior and posterior cortical regions. This indicates that regionally structured but broadly distributed vasodynamic propagation patterns are a generic feature across subjects

**Figure 6.**
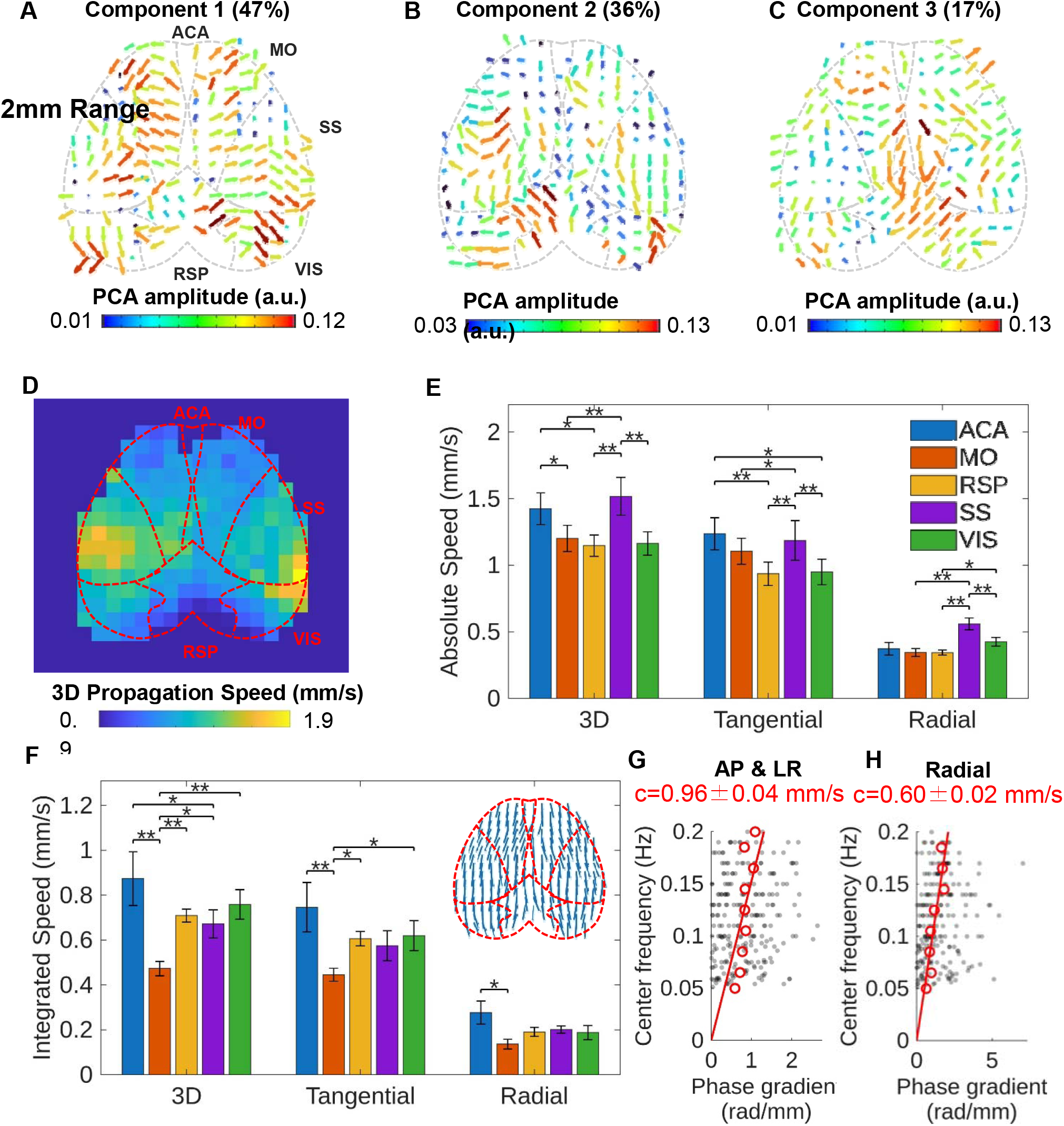
Group-level spatial pattern of 3D vasodynamic propagation speed. (A–C) The top three principal components from PCA of the 3D propagation speed maps across all 118 trials at a 2 mm fitting window, explaining 47% (A), 36% (B), and 17% (C) of total variance, respectively. Arrow length reflects the tangential projection of the vector magnitude; color encodes PCA amplitude. Cortical region boundaries are shown as dashed lines. (D) Averaged 3D propagation speed amplitude retained at positions passing a one-sample t-test (p < 0.01). (E) Absolute propagation speed (median of per-voxel magnitudes) for the 3D, tangential, and radial components across five cortical ROIs (ACA, MO, RSP, SS, and VIS). SS exhibited significantly higher absolute speed (repeated-measures ANOVA with Bonferroni post-hoc correction, *p < 0.05, **p < 0.01). (F) Integrated propagation speed (magnitude of the mean signed velocity vector across voxels) for the same three components and ROIs. ACA exhibited the highest integrated speed and MO the lowest (repeated-measures ANOVA with Bonferroni post-hoc correction, *p < 0.05, **p < 0.01). The averaged tangential propagation direction map shows a consistent anterior-to-posterior tendency (Rayleigh test on doubled angle, p < 0.01). Error bars in (E) and (F) represent standard error across animals (n = 18). (G) Scatter plot shows tangential (AP and LR) phase gradients from all 118 trials of 8Hz cortical CBV fMRI in a 1.8x1.8 mm SS ROI, as a function of center frequency, with the estimated propagation speed c = 0.96 ± 0.04 mm/s, R2=0.54 obtained by linear regression. (H) Scatter plot shows radial phase gradients from all 118 trials of 8Hz cortical CBV fMRI in a 2.2 x 2.2 mm SS ROI, as a function of center frequency, with the estimated propagation speed c = 0.60 ± 0.02 mm/s, R2=0.81 obtained by linear regression.

As a complement to the PCA analysis, we quantified propagation dynamics by computing the trial-averaged propagation speed along the tangential and radial axes, as well as the integrated 3D vector magnitude (Fig. 6D-H). Tangential propagation across the cortex presented mean speeds ranging from approximately 0.8 to 1.5 mm/s, exhibiting spatial variability across cortical regions with a structured distribution along the anterior–posterior axis (Supp. Fig. 6B, Fig. 6F). In contrast, radial propagation speed was substantially smaller at approximately 0.25 to 0.65 mm/s (Supp. Fig. 6C), consistent with the more heterogeneous and depth-dependent nature of vasodynamic timing. Regions exhibiting higher radial speeds were sparsely distributed and concentrated preferentially at the lateral cortical margins, where the cortical mantle in the flattened geometry is thinnest (Supp. Fig. 4) and the radial estimate is more influenced by superficial vasculature, including major pial arterial branches (Supp. Fig. 6C). Such superficial vascular contributions could compress phase differences across cortical depth, yielding shallower local radial gradients and consequently higher radial speed estimates.

The 3D propagation speed, combining both tangential and radial components, ranged from approximately 0.9 to 1.9 mm/s across the cortex (Fig. 6D), capturing the net effect of these anisotropic propagation patterns. To quantify regional variation, propagation speeds were measured within five anatomically defined cortical ROIs, including the anterior cingulate area (ACA), somatomotor area (MO), somatosensory area (SS), visual area (VIS), and retrosplenial area (RSP). Propagation speed, measured as the median per-voxel magnitude within each ROI, was significantly highest in SS across 3D and radial components, whereas ACA showed the highest mean tangential speed (repeated-measures ANOVA with Bonferroni correction, p < 0.01; Fig. 6E), indicating component-specific regional variation in local vasodynamic propagation speed. To assess how consistently propagation is organized within each region, we computed the integrated speed as the magnitude of the mean signed velocity vector across voxels. The averaged tangential direction map showed a consistent anterior-to-posterior tendency across cortex (Rayleigh test on doubled angle, p < 0.01; Fig. 6F), and the regional integrated speed reflected the degree of coherent vasodynamic propagation within each ROI. The ACA exhibited the highest integrated speed and MO the lowest across 3D, tangential, and radial components (Fig. 6F), indicating that propagation within ACA is directionally coherent, whereas MO, situated at the boundary between anterior and middle cerebral artery territories^51,52^, receives more mixed directional inputs that partially cancel in the signed average. To further validate the propagation speed range, median phase gradients were regressed against center frequency across all 118 trials, yielding c = 0.96 ± 0.04 mm/s (R² = 0.54) along the tangential axes (Fig. 6G) and c = 0.60 ± 0.02 mm/s (R² = 0.81) along the radial axis (Fig. 6H), which are consistent with the ROI-level speeds and confirmed the anisotropic organization described above. Cortical areas therefore differ in both the magnitude and the directional coherence of vasodynamic propagation, and the frequency-resolved regression independently reproduces the ROI-level speed estimates. Across the full dataset, these analyses resolve spontaneous cortical vasodynamics as vessel-aligned, frequency-specific oscillations that propagate with an anisotropic and regionally organized structure across the awake cortex.

## Discussion

In this work, we developed a cross-scale framework for mapping vasodynamic propagation across the cortex in the awake mouse with CBV-fMRI at 14 T. We showed that vasodynamic activity exhibited coordinated organization across spatial scales, from vessel-associated oscillations (Fig. 2) to mesoscopic propagation patterns and cortex-wide structured dynamics (Fig. 3, 5). Using space-frequency SVD analysis, the frequency-specific phase gradients were estimated in both cortex-wide tangential plane and laminar-specific radial axis to build a 3D propagation speed map, presenting region-specific propagation patterns of vasodynamics (Fig. 5, 6). Validation using a bilateral forepaw stimulation paradigm with a fixed interhemispheric delay confirmed that SF-SVD accurately recovers phase offsets between cortical regions (Fig. 4), and application to independent fUS data reproduced comparable frequency-specific coherence and phase gradient patterns (Supp. Fig. 3), indicating that the observed propagation structure reflects an intrinsic property of cortical vasodynamics rather than a modality-specific artifact. Together, these results establish CBV-fMRI-based SF-SVD as a noninvasive approach for resolving multiscale vasodynamic propagation across the cortex under awake conditions.

### Toward Vessel-Resolved Mapping of Vasodynamics Using CBV-Weighted Resting-State fMRI

Resting-state fMRI studies have long demonstrated that spontaneous brain activity exhibits structured organization across both spatial and frequency domains, forming the basis of functional connectivity analysis^53,54^. Subsequent work has shown that these signals can be decomposed into frequency-specific components using predefined sub-bands, time–frequency coherence analyses, and multiscale spectral methods, revealing frequency-dependent variations in network organization and physiological contributions^55–59^. In parallel, spatiotemporal analyses have identified structured propagation and lag relationships across cortical networks, suggesting that resting-state activity is organized as traveling waves with hierarchical dynamics^14,60^. ICA and related data-driven approaches have been widely used to separate these fluctuations into spatially structured components, including neural networks and physiological signals that are typically classified as noise and removed during preprocessing^61–64^. While these developments establish a rich framework for analyzing the spectral and spatial organization of resting-state signals, they are largely based on BOLD contrast, in which signal fluctuations reflect a composite of vascular, metabolic, and neuronal processes. This intrinsic mixing limits the ability to directly attribute frequency-specific structure to underlying vasodynamics, and vascular contributions are therefore typically modeled as confounds rather than interrogated as an organized physiological system.

CBV-weighted fMRI provides a critical complement to other modalities of fMRI because it offers improved vascular specificity while preserving recognizable resting-state network structure. Prior studies using VASO and contrast-enhanced CBV imaging have demonstrated that canonical functional networks can be robustly identified using ICA, confirming that CBV signals preserve large-scale functional organization while offering enhanced spatial specificity ^38,65–67^. However, these approaches have largely focused on network-level organization or bulk signal properties, without resolving the vascular origin or spatial organization of vasodynamic oscillations. Building on prior single-vessel CBV/BOLD fMRI work showing vessel-specific ultra-slow fluctuations and their spatial structure^18,41,42^, the present study extends this framework to the awake resting brain and shows that spontaneous vasodynamic activity can be resolved as vessel-aligned, frequency-specific spatial modes across the cortex. In Figs. 1 and 2, ICA analysis first isolated a dominant slow component aligned with penetrating vessels, whereas SF-SVD further revealed that these broadband vascular signals (0.01-0.2 Hz) were in fact composed of multiple vessel-specific oscillatory modes distributed across frequency.

The spatial organization was refined by NMF analysis of the vessel-specific coherence magnitude maps. Cortical components were preferentially weighted in the 0.06-0.2 Hz range, whereas the low band below 0.06 Hz included a relatively stronger subcortical contribution (Fig. 3D, E). Although prior resting-state studies did not resolve vasodynamics with vessel specificity, several observations are consistent with this cortical/subcortical spectral stratification. The prior two-photon work in awake mice detected cortical arteriolar vasomotion prominently across the 0.06-0.2 Hz regime^17^, including long-wavelength traveling waves along pial and penetrating arterioles. By contrast, resting-state and arousal-related fMRI studies have repeatedly implicated thalamic and basal-forebrain systems in very slow fluctuations that shape large-scale spontaneous activity underlying arousal state changes^68–70^. More broadly, whole-brain spectral analyses of resting-state fMRI have shown that cortical and subcortical regions do not contribute uniformly across frequency bands, even though those BOLD-based studies could not isolate the vascular component directly^68,71^. Our group-level CBV analysis extended cortical arteriole-specific vasomotion measurements to a brain-wide framework and identified a frequency-dependent cortical vasodynamic organization that provided the foundation for the 3D propagation mapping.

### Multiscale Origins of Vasodynamic Propagation: From Intrinsic Vasomotion to Neuromodulatory Control

The vasodynamic organization observed here is unlikely to reflect a single propagation mechanism acting uniformly across the cortex. Instead, the data are more consistent with intrinsic vasomotion as the dominant local source, with its observable phase structure shaped at larger scales by vascular compartmentation, laminar timing differences, and state-dependent neuromodulation. Prior single-vessel CBV-fMRI established that ultra-slow arteriole-dominated fluctuations can be resolved with vessel specificity and exhibit millimeter-scale spatial structure^18^, and awake optical imaging further showed that vasomotor phase evolves as traveling waves along pial and penetrating arterioles^17^. In particular, vasomotion-related propagation was quantified from both vessel diameter and smooth-muscle Ca²O signals, yielding phase gradients on the order of ∼0.3-0.4 rad/mm and mean speeds near ∼2 mm/s along the superficial arteriole network^17^. The present study extends that framework from individual vessels to mesoscopic and cortex-wide CBV-based mapping. In the high-resolution CBV-fMRI data, phase gradients estimated over ∼4 mm cortical regions were consistently ∼0.4-0.5 rad/mm, with a corresponding propagation speed of ∼2.3 mm/s (Supp. Fig. 2D); functional ultrasound yielded a closely matching gradient of ∼0.40 rad/mm using the same SF-SVD analysis (Supp. Fig. 3E). Together with the tangential-plane propagation speeds measured in the 8 Hz dataset, these results indicate that the tangential, vessel-aligned component of the CBV signal falls within the same regime as previously observed vasomotion traveling waves (Fig. 5), while extending those observations from optical surface measurements to cortex-wide mesoscopic mapping with a translational MRI-compatible modality.

By contrast with tangential propagation, the radial phase gradients resolved by the 8 Hz CBV-fMRI data point to a different regime. Significant radial gradients were sparse, bidirectional, and locally heterogeneous, and the corresponding radial propagation speeds were substantially slower, approximately 0.25-0.65 mm/s, whereas tangential propagation remained in the ∼0.8-1.5 mm/s range at the 2 mm fitting window and ∼1.8-2.8 mm/s at 4 mm (Supp. Fig. 6). Thus, at a comparable vasomotion frequency, the steeper radial phase gradients imply markedly slower effective propagation than the ∼2 mm/s along-vessel regime reported for pial and penetrating arterioles. This makes it unlikely that the radial gradients observed with CBV-fMRI simply reflect vasomotion traveling along single penetrating vessels. A more plausible interpretation is that they represent an emergent laminar timing structure, in which intrinsic vasomotion propagation can be integrated or even filtered through depth-dependent vascular compartments and local timing delays arising from neurovascular coupling and neuromodulatory regulation.

This interpretation helps reconcile three apparently different observations in the literature. First, cortical depth-specific optical measurements showed that microvascular dilation onset and peak timing vary systematically with depth, with deeper arteriolar elements responding earlier and superficial capillary compartments responding later; this is exactly the kind of built-in laminar timing structure that could generate radial phase offsets in spontaneous CBV signals^72^. Second, the long-wavelength vasomotion study emphasized that the ∼2 mm/s regime likely reflects the more faithful speed of a pure along-arteriole traveling wave and argued that shorter vessel segments can bias estimates toward steeper phase gradients and therefore slower inferred velocities^17^. Indeed, awake optical measurements of spontaneous pial vasomotion reported heterogenous propagation across short, i.e., 0.3-0.5 mm vessel length vessels with propagation speeds down to ∼0.4 mm/s^17,73^.

A further implication is that intrinsic vascular propagation itself is unlikely to be governed by a single conduction mode. TRPA1-dependent endothelial signaling shows that upstream vasodilatory responses can be initiated through a biphasic mechanism, with slow Ca²□-dependent propagation through capillary endothelium followed by conversion into faster electrical conduction in post-arteriole transitional segments and upstream arterioles^30,74^. This provides an intrinsic vascular mechanism by which slow and fast propagation regimes can coexist within the same network. Within this framework, the radial gradients observed here are physiologically feasible without requiring them to represent the same arteriole-wall mechanism measured optically along pial vessels. This broader laminar regulation could also be shaped by state-dependent modulatory systems. Noradrenergic and cholinergic pathways both regulate vascular tone^26,48^, exhibit nonuniform laminar organization^49,50^, impose different temporal structure on cortical state^75^, and modulate ultra-slow vascular dynamics^25,27^, making them plausible contributors to radial phase structure. The key point is therefore not that radial gradients arise from a mechanism unrelated to vasomotion, but that they likely reflect vasodynamic signals after the vasomotion has been filtered through additional layers of vascular and brain-state regulation.

### Multiscale Vasodynamic Organization as a Candidate Readout of Vascular Network Integrity

Group-level decomposition of trial-wise 3D propagation-speed maps revealed that vasodynamic propagation is not spatially uniform but instead organizes into partition-like modes across the cortex with a 2 mm fitting scale (Fig. 6A-C). These maps were derived from 200 µm isotropic voxels, with local phase gradients fit over a 2 mm tangential window and a radial window of at least 0.8 mm, and then resampled onto a 0.4 mm grid for decomposition. The spatial scale of the propagation estimate is therefore set by the fitting window rather than by the acquisition voxel size. These dominant components form spatially structured patterns with distinct medial–lateral and anterior–posterior propagation tendencies across cortical territories, indicating that vasodynamic propagation is regionally organized rather than uniformly distributed. When the fitting window was expanded to 4 mm, the corresponding components became visibly smoother and more axis-aligned (Supp. Fig. 7), indicating that the analysis shifts from finer-scale propagation structure toward larger-scale vasodynamic organization. This larger-scale vasodynamic organization is supported by the persistent anterior-posterior directional tendency in the tangential angle map (Fig. 6F, Rayleigh test p < 0.01) and by the known topology of the mouse pial arterial network, in which interconnected ACA-, MCA-, and PCA-supplied surface routes distribute flow across the cortex before feeding penetrating arterioles^51,52^. The anterior-posterior tendency is also consistent with the dominant direction reported for cortical-surface vasomotion waves in prior optical imaging work^17^. Together, these findings indicate that vasodynamic propagation is organized across multiple spatial scales, from finer local structure to broader cortex-wide domains. This organization parallels resting-state fMRI observations of reproducible partitions, gradients, and lag-defined propagation motifs^14,60,76–78^, and suggests that spontaneous vasodynamics can provide a vascular organizing framework for large-scale brain mapping.

This multiscale structure of cortical vasodynamic propagation also raises a translational possibility. Since the propagation structure reflects the coordinated function of arteriolar smooth muscle, endothelial conduction, neurovascular coupling, and neuromodulatory vascular control, disorders that disrupt these processes could plausibly alter key features of vasodynamic propagation itself. In the present data, this organization is captured both by the partition-like component maps and by spatially structured tangential, radial, and integrated propagation-speed measures across cortex (Fig. 6, Supp. Fig. 7). These region-specific vasodynamic speed measurements raise the possibility that diseased brains may be characterized not simply by weaker vascular oscillations, but by altered propagation topology, directional bias, or laminar timing structure. The larger-scale vasodynamic motifs may serve as candidate readouts of distributed vascular dysfunction in disorders such as cerebral amyloid angiopathy, cerebral small vessel disease, and vascular cognitive impairment, where abnormalities in vasomotion, neurovascular coupling, and cerebrovascular reactivity are already implicated^22,26,79,80^. Existing human MRI biomarkers, including cerebrovascular reactivity, low-frequency hemodynamic measures, and combined resting-state fMRI/ASL indices of neurovascular dysfunction, capture related aspects of vascular physiology, but do not directly assess how vascular timing is organized across the cortical sheet^81–83^. Human CBV-sensitive fMRI is feasible with ferumoxytol- and VASO-based approaches^38,84,85^, and individual human arteries, from large feeding vessels to cortical arteries, can be measured without exogenous agents using phase-contrast functional angiography and vessel-scale fMRI^86,87^. Given these capabilities, the broader propagation motifs resolved here at larger spatial scales may offer a plausible translational target, although this remains a hypothesis that will require direct validation in disease models and human cohorts.

### Limitations

Several limitations warrant consideration. First, although the near-100-µm CBV-fMRI resolution used here is sufficient to anatomically identify penetrating vessels, the measured vasodynamic signal within each voxel still reflects an integrated intravoxel vascular readout rather than a truly vessel-segment-specific measurement as in optical imaging. The present approach therefore represents a shift from vessel-visible anatomy to mesoscale vasodynamic mapping. This distinction is important for interpreting the phase-gradient results that finer-scale estimates are more likely to reflect mixed local regulatory processes, whereas larger-scale estimates are more likely to be dominated by propagation along larger arteries and thus by classical vasomotion. Second, temporal sampling imposes an additional constraint. The 2 s TR of the high-resolution CBV-fMRI data limits the minimum distance over which phase gradients can be robustly estimated and reduces sensitivity to faster or shorter-range propagation structure. Although the vessel-specific coherence patterns suggest that the measured vasodynamics are not primarily driven by non-physiological motion artifacts, the fMRI data remain susceptible to temporal aliasing. To address this limitation, we implemented 8 Hz sampling to improve sensitivity to short-distance phase gradients and reduce temporal aliasing of physiological fluctuations^2,88^. Finally, although we observed markedly different propagation speeds in the tangential plane and along the radial axis, the present measurements still capture only the integrated vasodynamic outcome of multiple underlying mechanisms. The current method cannot disentangle the relative contributions of intrinsic vasomotion, conducted vascular signaling, local neurovascular coupling, capillary-level regulation, or neuromodulatory coordination. Disentangling these regulatory sources will be essential before vasodynamic propagation can be interpreted confidently as a translational biomarker.

## Materials and Methods

### Animal Procedures

A total of 29 C57BL/6J wild-type mice (14 male, 15 female, 6-12 months old) were used for the fMRI experiments. Mice were group-housed (2-4 per cage) under a 12-hour light/dark cycle with food and water ad libitum. All animal procedures were conducted in accordance with protocols approved by the Massachusetts General Hospital Institutional Animal Care and Use Committee, and animals were cared for according to the requirements of the National Research Council’s Guide for the Care and Use of Laboratory Animals. Some animals contributed to more than one imaging protocol.

All mice underwent surgery under anesthesia (1-2% isoflurane with 1 L/min medical air) to implant an RF coil. Briefly, anesthetized mouse heads were stabilized in a stereotaxic stage, and the skull was exposed after aseptic preparation. A 600 MHz single-loop surface RF coil was secured to the skull with cyanoacrylate glue and dental cement to ensure stability. Analgesic (buprenorphine, 0.1-0.3 mg/kg) was administered before the procedure and periodically for three days post-surgery. Mice were allowed at least one week of recovery before acclimation began.

Following recovery, animals were acclimated to the MRI environment using a previously published head-fixation training protocol^47^. Briefly, mice were scanned twice weekly for approximately one month, yielding 2-4 trials per scan session. For all CBV scans, mice were injected with MION particles (25 mg/kg) via intraperitoneal injection approximately 60 minutes before scanning to allow for stable blood-pool distribution.

For the phase validation experiment, bilateral forepaw electrical stimulation was delivered through subcutaneous needle electrodes inserted into each forepaw. Rectangular current pulses (1 mA amplitude, 0.5 ms duration, 8 Hz pulse train) were generated by an isolated pulse stimulator. The paradigm consisted of alternating 40-second cycles, each composed of an 8-second stimulation period followed by a 32-second rest period. Right forepaw stimulation began at cycle onset, and left forepaw stimulation was delayed by 5 seconds relative to the right, creating a predictable interhemispheric timing offset at 0.025 Hz. A total of 8 mice were used for this experiment.

### MRI Acquisition

MRI data were acquired with a 14 T horizontal MRI scanner (Magnex Scientific, UK) located at the Athinoula A. Martinos Center for Biomedical Imaging in Boston, MA. The magnet is equipped with a Bruker Avance Neo console (Bruker BioSpin, Billerica, MA) and operated using ParaVision 360 V3.3. A microimaging gradient system (Resonance Research Inc., Billerica, MA) provides a peak gradient strength of 1.2 T/m over a 60-mm diameter.

For the high-resolution whole-brain protocol, anatomical images were obtained with a multi-slice T1-weighted 2D gradient echo FLASH sequence (TE: 2.7 ms, TR: 1250 ms, flip angle: 90°, 4 averages, 100×100×200 µm resolution). Functional CBV images were acquired using a multi-slice 2D EPI sequence (TE: 6.2 ms, TR: 1 s, two segments, effective volume TR = 2 s; bandwidth: 278 kHz, 100×100 µm in-plane resolution, 200 µm slice thickness). Each trial consisted of 205 repetitions for a total duration of 410 s.

For the cortex-specific ultra-fast protocol, two T1-weighted 2D gradient echo FLASH anatomical scans were acquired (TE: 2.7 ms, TR: 500 ms, flip angle: 70°, 2 averages, 200×200×200 µm resolution): one with coverage matching the functional EPI and one whole-brain scan with an increased number of slices for registration. Functional CBV images were acquired using a 2D EPI sequence at 8 Hz sampling (TE: 4 ms, TR: 0.125 s, bandwidth: 250 kHz, 200×200×200 µm isotropic resolution). Each trial consisted of 3280 repetitions for a total duration of 410 s.

### MRI Data Preprocessing

The fMRI images were processed using the Analysis of Functional Neuroimages (AFNI) software^89^. For the high-resolution EPI datasets, the preprocessed data from a standard AFNI pipeline^36,47^ were used, which included registration to the Australian Mouse Brain Mapping Consortium (AMBMC) template, despiking (3dDespike), motion correction (3dvolreg), and brain masking using the AMBMC template. For the cortex-specific ultra-fast scan data, a custom registration workflow was used. The functional EPIs were first registered to the co-acquired cortex-specific FLASH anatomical image. This cortex FLASH was then registered to the whole-brain FLASH, which was in turn registered to the AMBMC template. The transformation matrices from the cortex-to-whole-brain and whole-brain-to-AMBMC registrations were combined and applied to the functional EPIs to achieve final template registration. Other preprocessing steps, including despiking, motion correction, and brain masking, were performed as for the high-resolution pipeline. Cortical regions were segmented based on the Allen Mouse Brain Atlas for subsequent region-specific analyses.

### Functional Ultrasound Imaging Animal Procedures and Acquisition

fUS experiments were performed in four C57BL/6J mice, including 2 males and 2 females at 1-3 months of age. Animals were group-housed under controlled environmental conditions on a 12-hour light/dark cycle with ad libitum access to food and water. All experiments were conducted in accordance with US National Institutes of Health guidelines and were approved by the Broad Institute of MIT and Harvard Institutional Animal Care and Use Committee.

Resting-state transcranial functional ultrasound imaging was acquired under light sedation using isoflurane and dexmedetomidine. Mice were induced with 4% isoflurane in oxygen (1.0 L/min), the scalp was shaved, and animals were positioned in a stereotaxic frame. During skin preparation and probe placement, anesthesia was maintained at approximately 2% isoflurane. A depilatory cream was applied briefly (less than 1 min) to ensure complete hair removal, and the scalp was thoroughly rinsed. Dexmedetomidine (0.05 mg/kg, s.c.) was then administered, and isoflurane was reduced to 0.5% in oxygen (0.5 L/min) for imaging. Data acquisition began at least 5 min after reaching 0.5% isoflurane. Physiological status was continuously monitored throughout imaging, including oxygen saturation (maintained at ≥95%) and heart rate (∼250-350 bpm).

Functional ultrasound imaging was performed using an Iconeus One scanner (Iconeus, Paris, France) equipped with an IcoPrime probe (15 MHz central frequency, 128 elements). The probe was positioned on the depilated scalp over the dorsal surface, and each session began with an angiographic scan spanning +3.0 to -3.0 mm relative to bregma (0.1 mm inter-slice spacing) to verify probe positioning and provide a vascular reference. Resting-state acquisitions were performed by sequentially imaging six coronal planes spanning +2.0 to -3.0 mm relative to bregma with 1.0 mm spacing between planes. For each plane, approximately 15 min of continuous data were acquired to obtain stable resting-state time series (effective TR approximately 3.6 s for the six-slice cycle). Slice positions were kept consistent across animals to enable standardized region-of-interest comparisons.

### ICA Analysis

ICA was applied to the high-resolution whole-brain CBV fMRI data using GIFT with automatic dimensionality estimation. The dominant component was selected as the one with the highest spatial overlap with the cortical vasculature identified in the CBV contrast image. The power spectral density of the component time course was estimated using the Welch method with a 512-sample window and 50% overlap.

### Space-Frequency SVD Analysis

Space-frequency SVD (SF-SVD) was applied following the spectral framework as shown in Supp. Fig. 1. The fMRI time series from all in-mask voxels were organized into a space-time data matrix X of dimension m × n, where m denotes the number of voxels and n the number of time points. Multitaper spectral estimation was performed using k = 7 Slepian tapers with a time-bandwidth product NW = 4. The space-frequency representation at each frequency bin *f* was computed by projecting each taper onto the Fourier basis, yielding a complex-valued m × k matrix. SVD was then applied to this matrix, producing a dominant left singular vector u (m × 1, complex-valued) and corresponding singular value *λ*. Coherence at each frequency was defined as *λ*₁² / ∑*_taper_ λ*², quantifying the fractional spectral power captured by the leading mode. The magnitude of u encodes the spatial distribution of coherent oscillatory power, and the angle of u encodes the relative phase of the oscillation across voxels. Here, the increasing phase corresponds to increasing lag time. The center frequencies of each trial were identified as the frequency with peak coherence above the baseline (>0.2 Hz) of more than one standard deviation, which were also above the 1/k noise floor (Supp. Fig. 8).

### Non-Negative Matrix Factorization

NMF was applied to the SF-SVD magnitude maps from all trials. Magnitude maps were vectorized into a matrix of dimension *n_trails_* x *m_voxels_* and z-score normalized across voxels. NMF with k = 10 components was computed using MATLAB’s nnmf function with a maximum of 50 iterations. Three primary components were selected for display based on their correspondence to anatomically interpretable cortical and subcortical regions from the Allen Mouse Brain Atlas. For each component, trial weights were grouped into center frequency bins (0.01-0.06, 0.06-0.135, and 0.135-0.2 Hz) and the mean weight ± standard error was computed across trials within each bin. Differences in the subcortical component weight across the three frequency bands were assessed with one-way ANOVA and Tukey-Kramer post-hoc testing.

### Phase Gradient Estimation from Cortical Regions of Interest

For the high-resolution whole-brain fMRI and fUS datasets, where the minimum resolvable phase difference exceeds 4 mm, phase gradients were estimated from approximately 4 mm cortical regions of interest covering multiple penetrating vessels visible in the SF-SVD magnitude map. Within each region, a linear regression was applied to the averaged phase as a function of distance from the region center. The slope of this regression gives the phase gradient magnitude |*g*| in rad/mm. Propagation speed was estimated by regressing the median phase gradient against center frequency across all trials, using the relation *c* = 2π*f* / |*g*|.

### Cortex Flattening

To separate propagation components tangential to and radial from the ultra-fast cortical fMRI, the curved cortex area was geometrically flattened. The three-dimensional coordinates of all in-mask cortical voxels were used to fit a second-order polynomial surface *f*(*x*, *y*) = *p_00_ + p₁₀x + p₀₁y + p₂₀x² + p₁₁xy + p₀₂y²* by least-squares minimization. The radial position of each voxel was redefined as its signed residual *z*’ = *z* — *f*(*x*, *y*), and the two tangential axes (anterior-posterior and medial-lateral) were reassigned based on arc-length distance along the fitted surface. After transformation, the depth axis at each voxel points perpendicular to the local cortical surface, and the two tangential axes lie within the cortical plane.

### Three-Dimensional Phase Gradient and Propagation Speed Estimation

For the 8 Hz cortical CBV fMRI dataset, voxel-wise 3D phase gradient vectors were estimated after cortex flattening. At each voxel, phase values of neighboring voxels within a sliding window along each of the three spatial axes were extracted using a tangential window of either 2 mm or 4 mm, depending on the analysis scale, and a radial window of at least 0.8 mm requiring at least 4 voxels through cortical depth. Linear regression was applied separately along each axis, and the gradient component was retained only when the regression reached statistical significance (p < 0.05 tangentially, p < 0.01 radially). The three components were assembled into a 3D vector at each voxel, and this vector was converted to a propagation speed vector by *c* = 2π*f* /*g* (applied element-wise along each axis). The resulting per-trial maps were spatially downsampled onto a 0.6 mm display grid by computing the vector median within each local ±0.6 mm neighborhood. To estimate frequency-resolved propagation speed, median phase gradients along tangential (AP and LR) and radial axes were regressed against center frequency across all 118 trials using the relation *c* = 2π*f* / |*g*| .

### PCA of Propagation Speed Maps

For group-level analysis, each 3D propagation speed map was resampled onto a 0.4 mm grid with a ±0.4 mm neighborhood to calculate the vector median. To characterize dominant spatial patterns across trials, the per-trial 3D propagation speed vectors from all cortical voxels were assembled into a matrix of dimension *n_trails_* x (*m_voxels_* x 3). Columns were normalized by median and median absolute deviation, compressed with an arctan transform (2/π) - *atan*(*x*) to reduce the influence of extreme values, and then z-scored. PCA was performed using a nan-aware pairwise covariance approach, in which each entry of the spatial covariance matrix was estimated from the trials with valid observations at both positions, a small ridge regularization was added to ensure positive semi-definiteness, and the leading eigenvectors were extracted. The top three principal components were visualized as spatial vector fields by projecting each component back onto the flattened cortical surface.

### Statistical Analysis

The group analysis of the propagation speed map was tested against zero using a one-sample t-test, and only positions reaching significance (p < 0.01) were retained for group-averaged maps. Directional consistency of tangential propagation was assessed by the Rayleigh test applied to doubled azimuth angles to account for the axial symmetry of the phase gradient (p < 0.01).

To compare propagation speed across cortical ROIs, per-trial speed values were averaged within each animal to obtain one measurement per ROI per animal (n = 18), capturing the consistent cross-animal spatial pattern. ROI differences were then tested using repeated-measures ANOVA with Bonferroni-corrected post-hoc pairwise comparisons, applied separately to the 3D, tangential, and radial components for both absolute and integrated propagation speed.

## Acknowledgements

This research was funded by NIH Brain Initiative funding (RF1NS124778, R01NS122904, R01NS120594), NSF grant 2123971, and the S10 instrument grant (S10 MH124733-01) to the Martinos Center.

## References

1 Uludağ, K. & Blinder, P. Linking brain vascular physiology to hemodynamic response in ultra-high field MRI. NeuroImage 168, 279–295 (2018). 10.1016/j.neuroimage.2017.02.063

2 Drew, P. J., Mateo, C., Turner, K. L., Yu, X. & Kleinfeld, D. Ultra-slow Oscillations in fMRI and Resting-State Connectivity: Neuronal and Vascular Contributions and Technical Confounds. Neuron 107, 782–804 (2020). 10.1016/j.neuron.2020.07.020

3 Zhou, X. A., Jiang, Y., Gomez-Cid, L. & Yu, X. Elucidating hemodynamics and neuro-glio-vascular signaling using rodent fMRI. Trends in Neurosciences 48, 227–241 (2025). 10.1016/j.tins.2024.12.010

4 Logothetis, N. K. What we can do and what we cannot do with fMRI. Nature 453, 869–878 (2008). 10.1038/nature06976

5 Hillman, E. M. C. Coupling Mechanism and Significance of the BOLD Signal: A Status Report. Annual Review of Neuroscience 37, 161–181 (2014). 10.1146/annurev-neuro-071013-014111

6 Kim, S. G. & Ogawa, S. Biophysical and physiological origins of blood oxygenation level-dependent fMRI signals. J Cereb Blood Flow Metab 32, 1188–1206 (2012). 10.1038/jcbfm.2012.23

7 Schaeffer, S. & Iadecola, C. Revisiting the neurovascular unit. Nature Neuroscience 24, 1198–1209 (2021). 10.1038/s41593-021-00904-7

8 Bolt, T. et al. Autonomic physiological coupling of the global fMRI signal. Nature Neuroscience 28, 1327–1335 (2025). 10.1038/s41593-025-01945-y

9 Hamel, E. Perivascular nerves and the regulation of cerebrovascular tone. J Appl Physiol (1985) 100, 1059–1064 (2006). 10.1152/japplphysiol.00954.2005

10 Mateo, C., Knutsen, P. M., Tsai, P. S., Shih, A. Y. & Kleinfeld, D. Entrainment of Arteriole Vasomotor Fluctuations by Neural Activity Is a Basis of Blood-Oxygenation-Level-Dependent &#x201c;Resting-State&#x201d; Connectivity. Neuron 96, 936–948.e933 (2017). 10.1016/j.neuron.2017.10.012

11 Intaglietta, M. Vasomotion and flowmotion: physiological mechanisms and clinical evidence. Vascular Medicine Review vmr-1, 101–112 (1990). 10.1177/1358836X9000100202

12 Buckner, R. L., Andrews-Hanna, J. R. & Schacter, D. L. The brain’s default network: anatomy, function, and relevance to disease. Ann N Y Acad Sci 1124, 1–38 (2008). 10.1196/annals.1440.011

13 Biswal, B. B. & Uddin, L. Q. The history and future of resting-state functional magnetic resonance imaging. Nature 641, 1121–1131 (2025). 10.1038/s41586-025-08953-9

14 Mitra, A., Snyder, A. Z., Hacker, C. D. & Raichle, M. E. Lag structure in resting-state fMRI. J Neurophysiol 111, 2374–2391 (2014). 10.1152/jn.00804.2013

15 Tong, Y. & Frederick, B. Tracking cerebral blood flow in BOLD fMRI using recursively generated regressors. Hum Brain Mapp 35, 5471–5485 (2014). 10.1002/hbm.22564

16 Raut, R. V. et al. Global waves synchronize the brain’s functional systems with fluctuating arousal. Sci Adv 7 (2021). 10.1126/sciadv.abf2709

17 Broggini, T. et al. Long-wavelength traveling waves of vasomotion modulate the perfusion of cortex. Neuron 112, 2349–2367.e2348 (2024). 10.1016/j.neuron.2024.04.034

18 He, Y. et al. Ultra-Slow Single-Vessel BOLD and CBV-Based fMRI Spatiotemporal Dynamics and Their Correlation with Neuronal Intracellular Calcium Signals. Neuron 97, 925–939.e925 (2018). 10.1016/j.neuron.2018.01.025

19 Iadecola, C. The Neurovascular Unit Coming of Age: A Journey through Neurovascular Coupling in Health and Disease. Neuron 96, 17–42 (2017). 10.1016/j.neuron.2017.07.030

20 Attwell, D. et al. Glial and neuronal control of brain blood flow. Nature 468, 232–243 (2010). 10.1038/nature09613

21 van Veluw, S. J. et al. Vasomotion as a Driving Force for Paravascular Clearance in the Awake Mouse Brain. Neuron 105, 549–561.e545 (2020). 10.1016/j.neuron.2019.10.033

22 Kozberg, M. G. et al. Vasomotion loss precedes impaired cerebrovascular reactivity and microbleeds in cerebral amyloid angiopathy. Brain Communications 7, fcaf186 (2025). 10.1093/braincomms/fcaf186

23 Di Marco, L. Y., Farkas, E., Martin, C., Venneri, A. & Frangi, A. F. Is Vasomotion in Cerebral Arteries Impaired in Alzheimer’s Disease? J Alzheimers Dis 46, 35–53 (2015). 10.3233/jad-142976

24 Fultz, N. E. et al. Coupled electrophysiological, hemodynamic, and cerebrospinal fluid oscillations in human sleep. Science 366, 628–631 (2019). doi:10.1126/science.aax5440

25 Hauglund, N. L. et al. Norepinephrine-mediated slow vasomotion drives glymphatic clearance during sleep. Cell 188, 606–622.e617 (2025). 10.1016/j.cell.2024.11.027

26 Korte, N. et al. Noradrenaline released from locus coeruleus axons contracts cerebral capillary pericytes via α2 adrenergic receptors. Journal of Cerebral Blood Flow & Metabolism 43, 1142–1152 (2023). 10.1177/0271678X231152549

27 Chuang, K.-H. et al. Cholinergic basal forebrain neurons regulate vascular dynamics and cerebrospinal fluid flux. Nature Communications 16, 5343 (2025). 10.1038/s41467-025-60812-3

28 Nilsson, H. & Aalkjaer, C. Vasomotion: mechanisms and physiological importance. Mol Interv 3, 79–89, 51 (2003). 10.1124/mi.3.2.79

29 Drew, P. J., Shih, A. Y. & Kleinfeld, D. Fluctuating and sensory-induced vasodynamics in rodent cortex extend arteriole capacity. Proceedings of the National Academy of Sciences 108, 8473–8478 (2011). 10.1073/pnas.1100428108

30 Longden, T. A. et al. Capillary K(+)-sensing initiates retrograde hyperpolarization to increase local cerebral blood flow. Nat Neurosci 20, 717–726 (2017). 10.1038/nn.4533

31 Mayhew, J. E. et al. Cerebral vasomotion: a 0.1-Hz oscillation in reflected light imaging of neural activity. Neuroimage 4, 183–193 (1996). 10.1006/nimg.1996.0069

32 Rayshubskiy, A. et al. Direct, intraoperative observation of ∼0.1Hz hemodynamic oscillations in awake human cortex: Implications for fMRI. NeuroImage 87, 323–331 (2014). 10.1016/j.neuroimage.2013.10.044

33 Earley, S., Kleinfeld, D. & van Veluw, S. J. Vasomotion of brain arteries: Ionic dynamics through systems physiology in health and vascular disease. Annual Review of Physiology (2027).

34 Schmitz, T. W. et al. Basal forebrain degeneration precedes and predicts the cortical spread of Alzheimer’s pathology. Nature Communications 7, 13249 (2016). 10.1038/ncomms13249

35 Weinshenker, D. Long Road to Ruin: Noradrenergic Dysfunction in Neurodegenerative Disease. Trends Neurosci 41, 211–223 (2018). 10.1016/j.tins.2018.01.010

36 Liu, X. et al. Identifying the bioimaging features of Alzheimer’s disease based on pupillary light response-driven brain-wide fMRI in awake mice. Nature Communications 15, 9657 (2024). 10.1038/s41467-024-53878-y

37 Kim, T. & Kim, S.-G. Cortical layer-dependent arterial blood volume changes: Improved spatial specificity relative to BOLD fMRI. NeuroImage 49, 1340–1349 (2010). 10.1016/j.neuroimage.2009.09.061

38 Huber, L. et al. High-Resolution CBV-fMRI Allows Mapping of Laminar Activity and Connectivity of Cortical Input and Output in Human M1. Neuron 96, 1253–1263.e1257 (2017). 10.1016/j.neuron.2017.11.005

39 Belliveau, J. W. et al. Functional mapping of the human visual cortex by magnetic resonance imaging. Science 254, 716–719 (1991). 10.1126/science.1948051

40 Lu, H. & van Zijl, P. C. A review of the development of Vascular-Space-Occupancy (VASO) fMRI. Neuroimage 62, 736–742 (2012). 10.1016/j.neuroimage.2012.01.013

41 Yu, X. et al. Sensory and optogenetically driven single-vessel fMRI. Nature Methods 13, 337–340 (2016). 10.1038/nmeth.3765

42 He, Y., Wang, M. & Yu, X. High spatiotemporal vessel-specific hemodynamic mapping with multi-echo single-vessel fMRI. J Cereb Blood Flow Metab 40, 2098–2114 (2020). 10.1177/0271678x19886240

43 Jiang, Y., Pais-Roldán, P., Pohmann, R. & Yu, X. High Spatiotemporal Resolution Radial Encoding Single-Vessel fMRI. Advanced Science 11, 2309218 (2024). 10.1002/advs.202309218

44 Mitra, P. P. & Pesaran, B. Analysis of dynamic brain imaging data. Biophys J 76, 691–708 (1999). 10.1016/s0006-3495(99)77236-x

45 Prechtl, J., Cohen, L., Pesaran, B., Mitra, P. & Kleinfeld, D. Visual stimuli induce waves of electrical activity in turtle cortex. Proceedings of the National Academy of Sciences 94, 7621–7626 (1997).

46 Kleinfeld, D. & Mitra, P. P. Spectral methods for functional brain imaging. Cold Spring Harb Protoc 2014, 248–262 (2014). 10.1101/pdb.top081075

47 Hike, D. et al. High-resolution awake mouse fMRI at 14 tesla. eLife 13, RP95528 (2025). 10.7554/eLife.95528

48 Lecrux, C. et al. Impact of Altered Cholinergic Tones on the Neurovascular Coupling Response to Whisker Stimulation. J Neurosci 37, 1518–1531 (2017). 10.1523/jneurosci.1784-16.2016

49 Morrison, J. H., Foote, S. L., O’Connor, D. & Bloom, F. E. Laminar, tangential and regional organization of the noradrenergic innervation of monkey cortex: dopamine-beta-hydroxylase immunohistochemistry. Brain Res Bull 9, 309–319 (1982). 10.1016/0361-9230(82)90144-7

50 Mechawar, N., Cozzari, C. & Descarries, L. Cholinergic innervation in adult rat cerebral cortex: a quantitative immunocytochemical description. J Comp Neurol 428, 305–318 (2000). 10.1002/1096-9861(20001211)428:2<305::aid-cne9>3.0.co;2-y

51 Blinder, P., Shih, A. Y., Rafie, C. & Kleinfeld, D. Topological basis for the robust distribution of blood to rodent neocortex. Proceedings of the National Academy of Sciences 107, 12670–12675 (2010). doi:10.1073/pnas.1007239107

52 Blinder, P. et al. The cortical angiome: an interconnected vascular network with noncolumnar patterns of blood flow. Nature Neuroscience 16, 889–897 (2013). 10.1038/nn.3426

53 Biswal, B., Yetkin, F. Z., Haughton, V. M. & Hyde, J. S. Functional connectivity in the motor cortex of resting human brain using echo-planar MRI. Magn Reson Med 34, 537–541 (1995). 10.1002/mrm.1910340409

54 Cordes, D. et al. Frequencies contributing to functional connectivity in the cerebral cortex in “resting-state” data. AJNR Am J Neuroradiol 22, 1326–1333 (2001).

55 Chang, C. & Glover, G. H. Time–frequency dynamics of resting-state brain connectivity measured with fMRI. NeuroImage 50, 81–98 (2010). 10.1016/j.neuroimage.2009.12.011

56 Yaesoubi, M., Allen, E. A., Miller, R. L. & Calhoun, V. D. Dynamic coherence analysis of resting fMRI data to jointly capture state-based phase, frequency, and time-domain information. Neuroimage 120, 133–142 (2015). 10.1016/j.neuroimage.2015.07.002

57 Yuen, N. H., Osachoff, N. & Chen, J. J. Intrinsic Frequencies of the Resting-State fMRI Signal: The Frequency Dependence of Functional Connectivity and the Effect of Mode Mixing. Frontiers in Neuroscience Volume 13 - 2019 (2019). 10.3389/fnins.2019.00900

58 Gohel, S. R. & Biswal, B. B. Functional integration between brain regions at rest occurs in multiple-frequency bands. Brain Connect 5, 23–34 (2015). 10.1089/brain.2013.0210

59 Cabral, J. et al. Cognitive performance in healthy older adults relates to spontaneous switching between states of functional connectivity during rest. Scientific Reports 7, 5135 (2017). 10.1038/s41598-017-05425-7

60 Raut, R. V., Snyder, A. Z. & Raichle, M. E. Hierarchical dynamics as a macroscopic organizing principle of the human brain. Proc Natl Acad Sci U S A 117, 20890–20897 (2020). 10.1073/pnas.2003383117

61 Beckmann, C. F., DeLuca, M., Devlin, J. T. & Smith, S. M. Investigations into resting-state connectivity using independent component analysis. Philos Trans R Soc Lond B Biol Sci 360, 1001–1013 (2005). 10.1098/rstb.2005.1634

62 McKeown, M. J. et al. Analysis of fMRI data by blind separation into independent spatial components. Human Brain Mapping 6, 160–188 (1998). 10.1002/(SICI)1097-0193(1998)6:3<160::AID-HBM5>3.0.CO;2-1

63 Griffanti, L. et al. ICA-based artefact removal and accelerated fMRI acquisition for improved resting state network imaging. Neuroimage 95, 232–247 (2014). 10.1016/j.neuroimage.2014.03.034

64. Kiviniemi, V. et al. Localization of the Resting State Vasomotor Fluctuation with FFT, Cross Correlation, Principal Component and Independent Component Analysis of fMRI data. Proc. Of International Society of Magnetic Resonance in Medicine 1708 (2001).

65 Magnuson, M., Majeed, W. & Keilholz, S. D. Functional connectivity in blood oxygenation level-dependent and cerebral blood volume-weighted resting state functional magnetic resonance imaging in the rat brain. J Magn Reson Imaging 32, 584–592 (2010). 10.1002/jmri.22295

66 Vanduffel, W. et al. Visual motion processing investigated using contrast agent-enhanced fMRI in awake behaving monkeys. Neuron 32, 565–577 (2001). 10.1016/s0896-6273(01)00502-5

67 Miao, X. et al. Detecting resting-state brain activity by spontaneous cerebral blood volume fluctuations using whole brain vascular space occupancy imaging. Neuroimage 84, 575–584 (2014).

68 Setzer, B. et al. A temporal sequence of thalamic activity unfolds at transitions in behavioral arousal state. Nature Communications 13, 5442 (2022). 10.1038/s41467-022-33010-8

69 Wang, M., He, Y., Sejnowski, T. J. & Yu, X. Brain-state dependent astrocytic Ca2+ signals are coupled to both positive and negative BOLD-fMRI signals. Proceedings of the National Academy of Sciences 115, E1647–E1656 (2018). 10.1073/pnas.1711692115

70 Chang, C. et al. Tracking brain arousal fluctuations with fMRI. Proceedings of the National Academy of Sciences 113, 4518–4523 (2016). 10.1073/pnas.1520613113

71 Zuo, X. N. et al. The oscillating brain: complex and reliable. Neuroimage 49, 1432–1445 (2010). 10.1016/j.neuroimage.2009.09.037

72 Tian, P. et al. Cortical depth-specific microvascular dilation underlies laminar differences in blood oxygenation level-dependent functional MRI signal. Proceedings of the National Academy of Sciences 107, 15246–15251 (2010). doi:10.1073/pnas.1006735107

73 Munting, L. P. et al. Spontaneous vasomotion propagates along pial arterioles in the awake mouse brain like stimulus-evoked vascular reactivity. J Cereb Blood Flow Metab 43, 1752–1763 (2023). 10.1177/0271678x231152550

74 Thakore, P. et al. Brain endothelial cell TRPA1 channels initiate neurovascular coupling. eLife 10, e63040 (2021). 10.7554/eLife.63040

75 Reimer, J. et al. Pupil fluctuations track rapid changes in adrenergic and cholinergic activity in cortex. Nature Communications 7, 13289 (2016). 10.1038/ncomms13289

76 Wig, G. S., Laumann, T. O. & Petersen, S. E. An approach for parcellating human cortical areas using resting-state correlations. Neuroimage 93 Pt 2, 276–291 (2014). 10.1016/j.neuroimage.2013.07.035

77 Gordon, E. M. et al. Generation and Evaluation of a Cortical Area Parcellation from Resting-State Correlations. Cereb Cortex 26, 288–303 (2016). 10.1093/cercor/bhu239

78 Gu, Y. et al. Brain Activity Fluctuations Propagate as Waves Traversing the Cortical Hierarchy. Cereb Cortex 31, 3986–4005 (2021). 10.1093/cercor/bhab064

79 Kisler, K. et al. Pericyte degeneration leads to neurovascular uncoupling and limits oxygen supply to brain. Nat Neurosci 20, 406–416 (2017). 10.1038/nn.4489

80 Choi, S. et al. Resting-state fMRI coherence is selectively diminished around 0.1 Hz in patients with unilateral carotid artery stenosis. NeuroImage, 121938 (2026). 10.1016/j.neuroimage.2026.121938

81 Dumas, A. et al. Functional magnetic resonance imaging detection of vascular reactivity in cerebral amyloid angiopathy. Ann Neurol 72, 76–81 (2012). 10.1002/ana.23566

82 Switzer, A. R. et al. Cerebrovascular reactivity in cerebral amyloid angiopathy, Alzheimer disease, and mild cognitive impairment. Neurology 95, e1333–e1340 (2020). 10.1212/wnl.0000000000010201

83 Li, H. et al. Dysfunction of neurovascular coupling in patients with cerebral small vessel disease: A combined resting-state fMRI and arterial spin labeling study. Experimental Gerontology 194, 112478 (2024). 10.1016/j.exger.2024.112478

84 D’Arceuil, H. et al. Ferumoxytol enhanced resting state fMRI and relative cerebral blood volume mapping in normal human brain. Neuroimage 83, 200–209 (2013). 10.1016/j.neuroimage.2013.06.066

85 Qiu, D., Zaharchuk, G., Christen, T., Ni, W. W. & Moseley, M. E. Contrast-enhanced functional blood volume imaging (CE-fBVI): enhanced sensitivity for brain activation in humans using the ultrasmall superparamagnetic iron oxide agent ferumoxytol. Neuroimage 62, 1726–1731 (2012). 10.1016/j.neuroimage.2012.05.010

86 Hu, Z. et al. Visual stimulus-evoked blood velocity responses in individual human posterior cerebral arteries measured with dynamic phase-contrast functional MR angiography. Imaging Neurosci (Camb) 3 (2025). 10.1162/IMAG.a.148

87 Proulx, S. et al. in Proceedings of the International Society for Magnetic Resonance in Medicine 0283 (International Society for Magnetic Resonance in Medicine, Honolulu, HI, USA, 2025).

88 Pais-Roldán, P., Biswal, B., Scheffler, K. & Yu, X. Identifying Respiration-Related Aliasing Artifacts in the Rodent Resting-State fMRI. Frontiers in Neuroscience Volume 12 - 2018 (2018). 10.3389/fnins.2018.00788

89 Cox, R. W. AFNI: software for analysis and visualization of functional magnetic resonance neuroimages. Comput Biomed Res 29, 162–173 (1996). 10.1006/cbmr.1996.0014

