## Supplemental Figures for "Spontaneous cortical vasodynamics form a multiscale propagation architecture across the awake brain"

**Supplementary Figures**

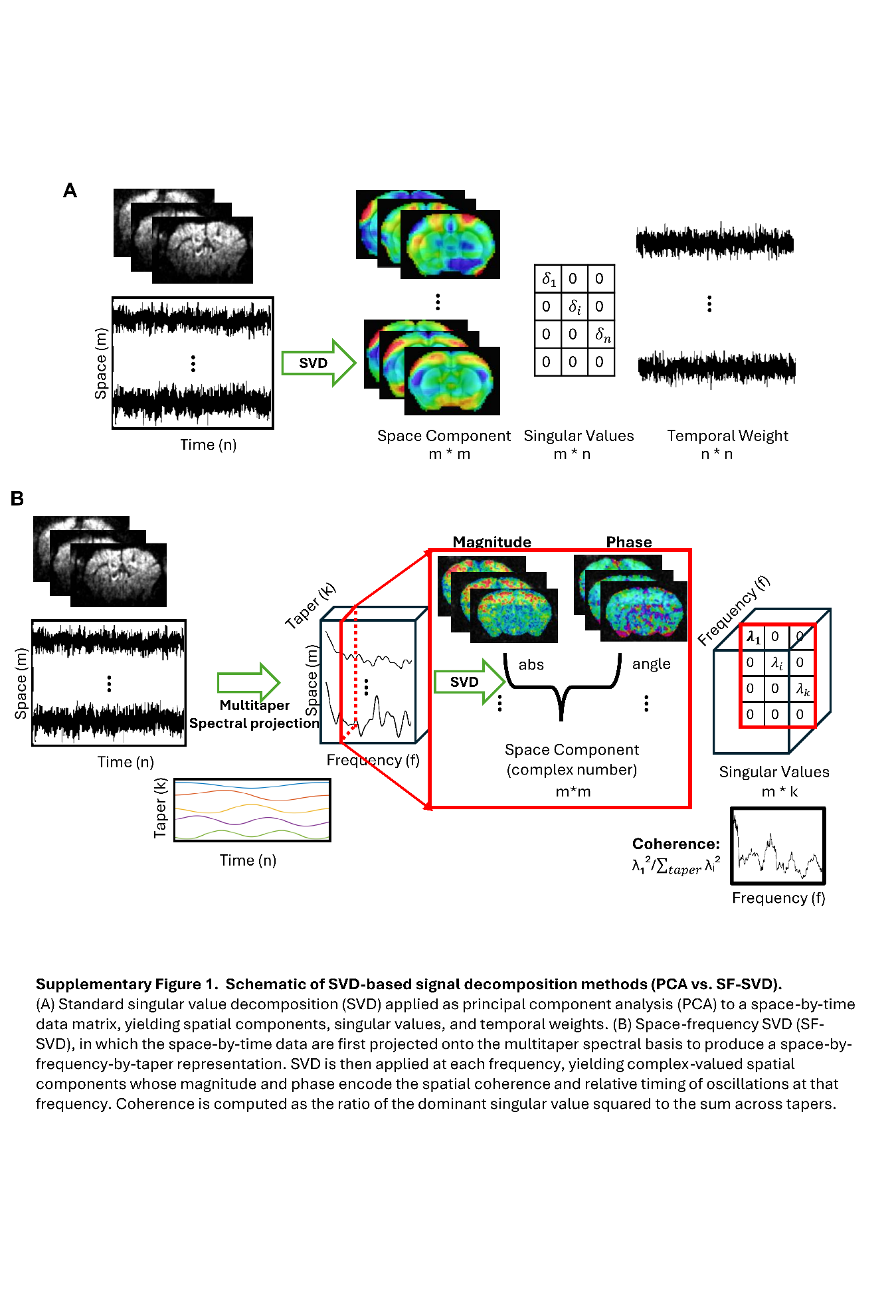

**Supplementary Figure 1. Schematic of SVD-based signal decomposition methods (PCA vs. SF-SVD).**

(A) Standard singular value decomposition (SVD) applied as principal component analysis (PCA) to a space-by-time data matrix, yielding spatial components, singular values, and temporal weights. (B) Space-frequency SVD (SF-SVD), in which the space-by-time data are first projected onto the multitaper spectral basis to produce a space-by-frequency-by-taper representation. SVD is then applied at each frequency, yielding complex-valued spatial components whose magnitude and phase encode the spatial coherence and relative timing of oscillations at that frequency. Coherence is computed as the ratio of the dominant singular value squared to the sum across tapers.

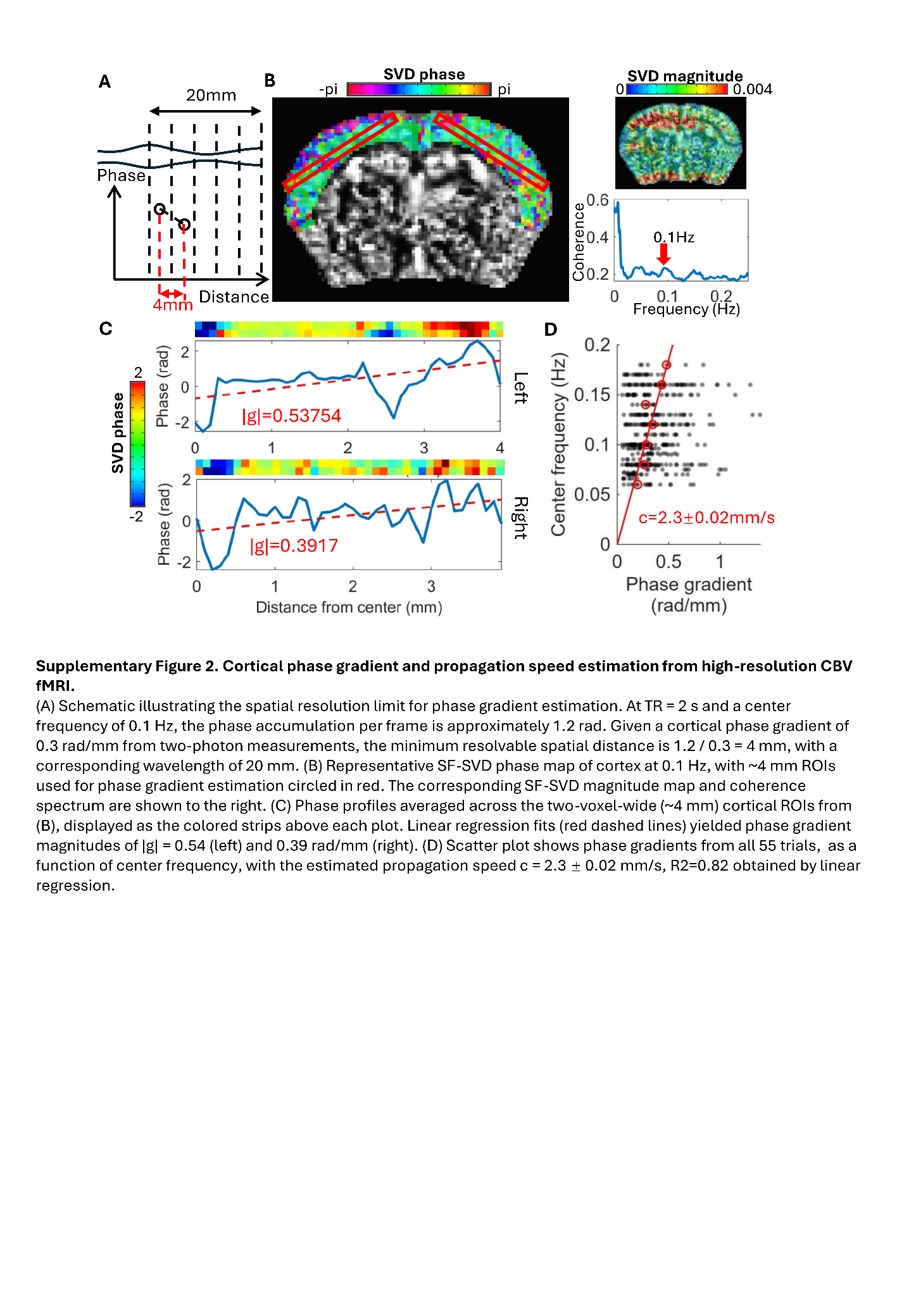

**Supplementary Figure 2. Cortical phase gradient and propagation speed estimation from high-resolution CBV fMRI.**

(A) Schematic illustrating the spatial resolution limit for phase gradient estimation. At TR = 2 s and a center frequency of 0.1 Hz, the phase accumulation per frame is approximately 1.2 rad. Given a cortical phase gradient of 0.3 rad/mm from two-photon measurements, the sampling-dependent minimum resolvable spatial distance is 1.2 / 0.3 = 4 mm, with a corresponding wavelength of 20 mm. (B) Representative SF-SVD phase map of cortex at 0.1 Hz, with ~4 mm ROIs used for phase gradient estimation circled in red. The corresponding SF-SVD magnitude map and coherence spectrum are shown to the right. (C) Phase profiles averaged across the two-voxel-wide (~4 mm) cortical ROIs from (B), displayed as the colored strips above each plot. Linear regression fits (red dashed lines) yielded phase gradient magnitudes of |g| = 0.54 (left) and 0.39 rad/mm (right). (D) Scatter plot shows phase gradients from all 55 trials, as a function of center frequency, with the estimated propagation speed c = 2.3 ± 0.02 mm/s, R2=0.82 obtained by linear regression.

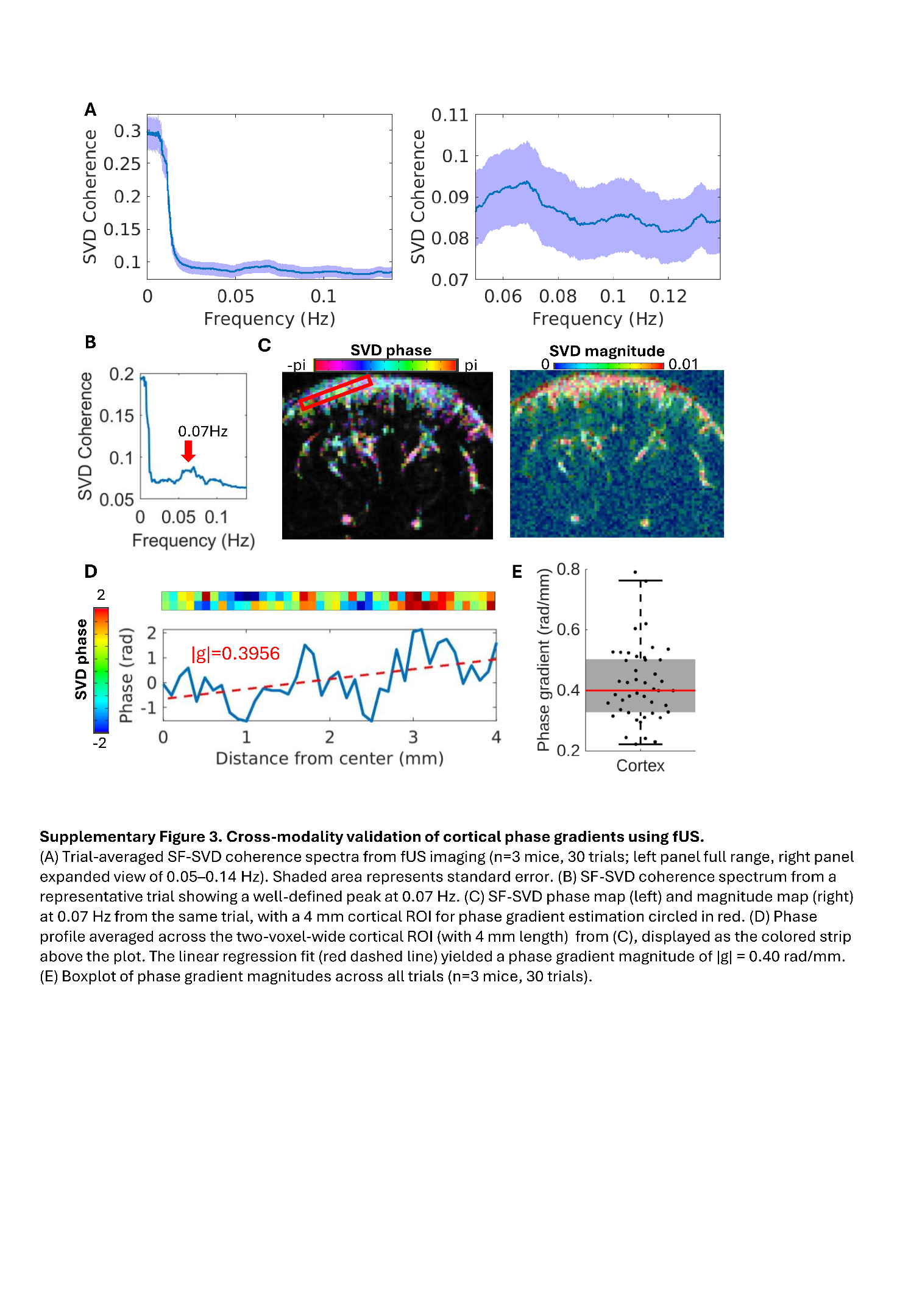

**Supplementary Figure 3. Cross-modality validation of cortical phase gradients using fUS.**

(A) Trial-averaged SF-SVD coherence spectra from fUS imaging (n=4 mice, 30 trials; left panel full range, right panel expanded view of 0.05–0.14 Hz). The shaded area represents standard error. (B) SF-SVD coherence spectrum from a representative trial showing a well-defined peak at 0.07 Hz. (C) SF-SVD phase map (left) and magnitude map (right) at 0.07 Hz from the same trial, with a 4 mm cortical ROI for phase gradient estimation circled in red. (D) Phase profile averaged across the two-voxel-wide cortical ROI (with 4 mm length) from (C), displayed as the colored strip above the plot. The linear regression fit (red dashed line) yielded a phase gradient magnitude of |g| = 0.40 rad/mm. (E) Boxplot of phase gradient magnitudes across all trials (n=4 mice, 30 trials).

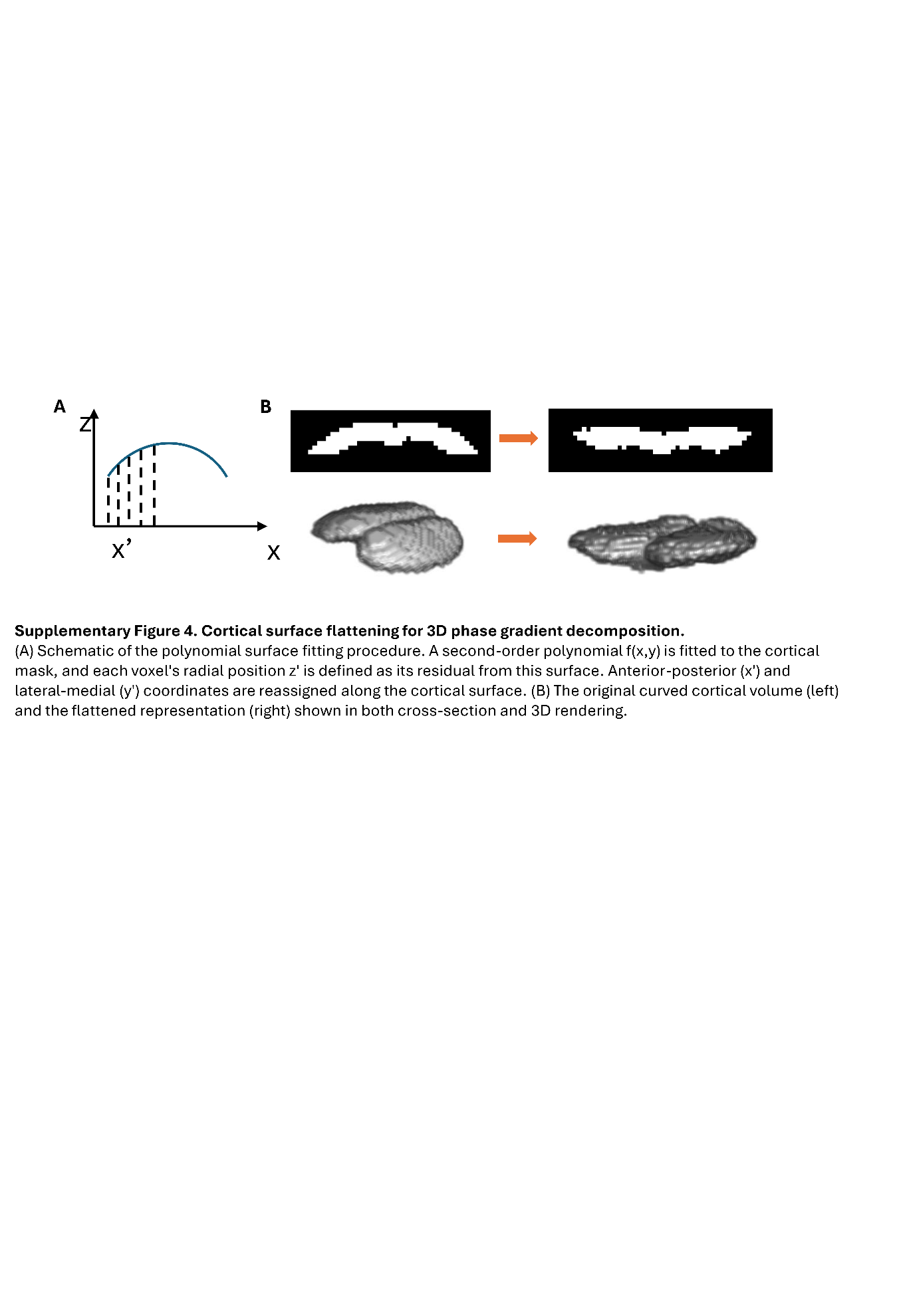

**Supplementary Figure 4. Cortical surface flattening for 3D phase gradient decomposition.**

(A) Schematic of the polynomial surface fitting procedure. A second-order polynomial f(x,y) is fitted to the cortical mask, and each voxel's radial position z' is defined as its residual from this surface. Anterior-posterior (x') and medial-lateral (y') coordinates are reassigned along the cortical surface. (B) The original curved cortical volume (left) and the flattened representation (right) shown in both cross-section and 3D rendering.

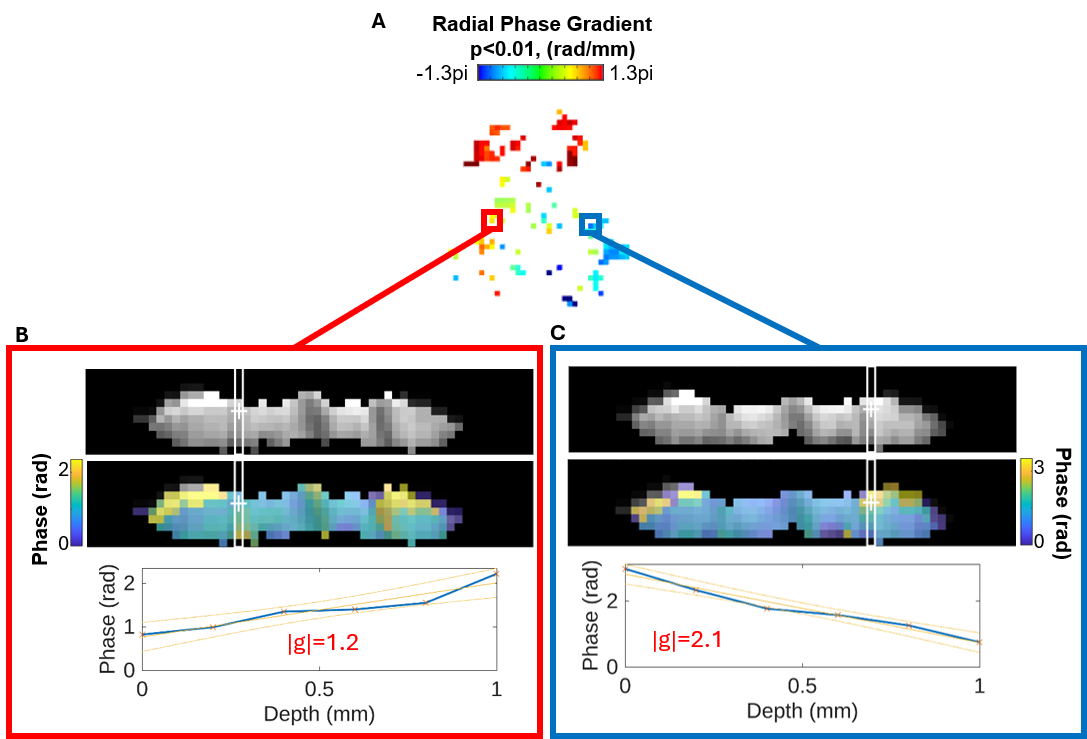

**Supplementary Figure 5. Representative bidirectional radial phase gradients at vessel-associated cortical sites.**

(A) Signed radial phase-gradient map from a representative 8-Hz CBV-fMRI trial at 0.11 Hz (Fig. 5F). Only cortical locations with significant linear phase-versus-depth slopes (p<0.01) are shown. Red and blue boxes denote representative sites with positive and negative radial phase gradients, respectively. (B–C) Flattened averaged EPI images (top), corresponding SF-SVD phase maps (middle), and radial phase profiles plotted as a function of relative cortical depth (bottom) for the selected sites. White outlines indicate the radial voxel columns included in the linear fits. The fitted gradients were |g|=1.2 rad/mm in (B) and |g|=2.1 rad/mm in (C). In both examples, the selected radial columns intersect dark, radially oriented features in the averaged EPI images, consistent with penetrating vessels. Markers show measured phase values, solid lines show linear fits, and dotted lines indicate 95% confidence intervals. Increasing phase corresponds to increasing lag; therefore, the opposing slopes indicate opposite directions of depth-dependent phase progression.

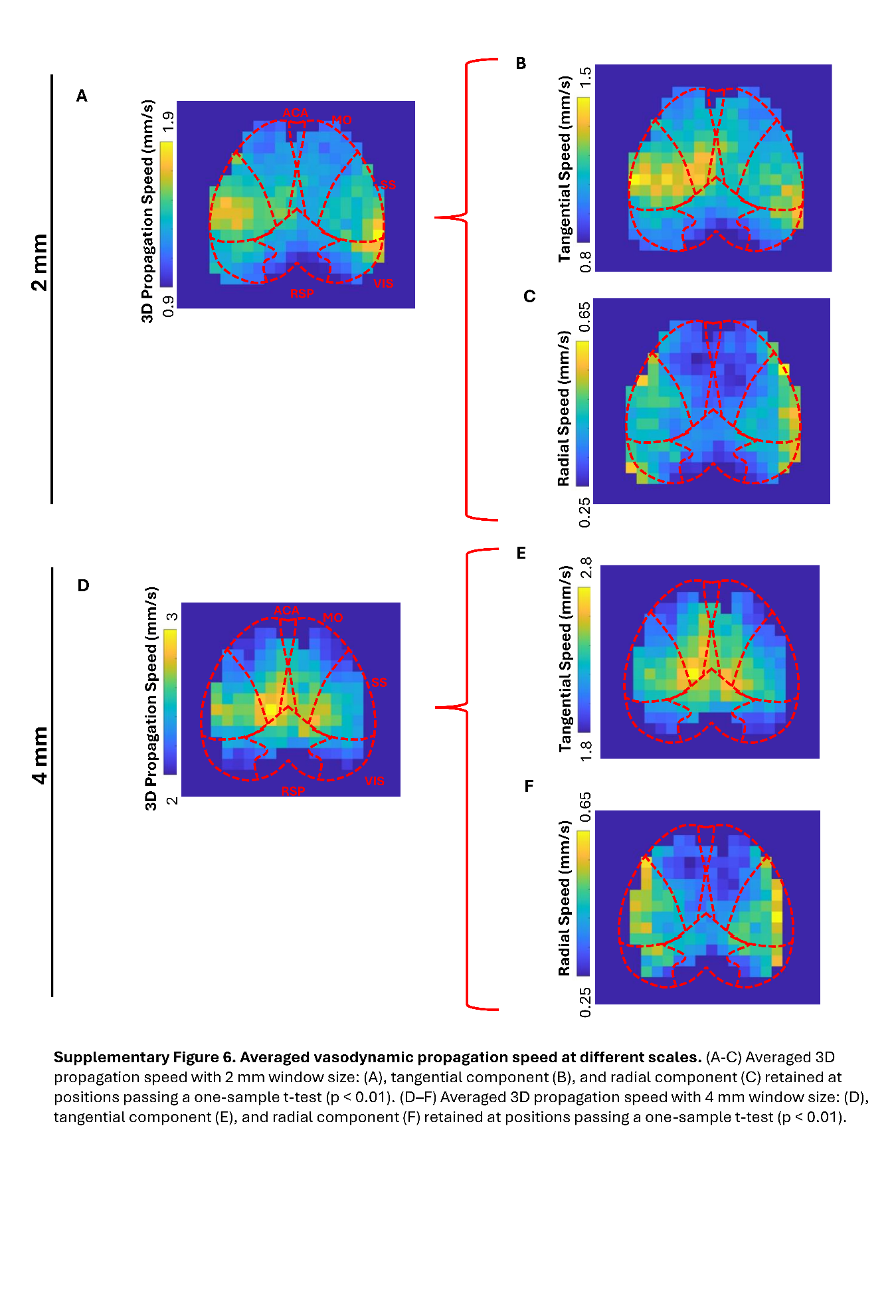

**Supplementary Figure 6. Averaged vasodynamic propagation speed at different scales.**

(A)–(C), 2 mm fitting window: (A) 3D propagation speed, (B) tangential component, and (C) radial component. (D)–(F), 4 mm fitting window: (D) 3D propagation speed, (E) tangential component, and (F) radial component. (one-sample t-test, p < 0.01, n=18 mice, 118 trials)

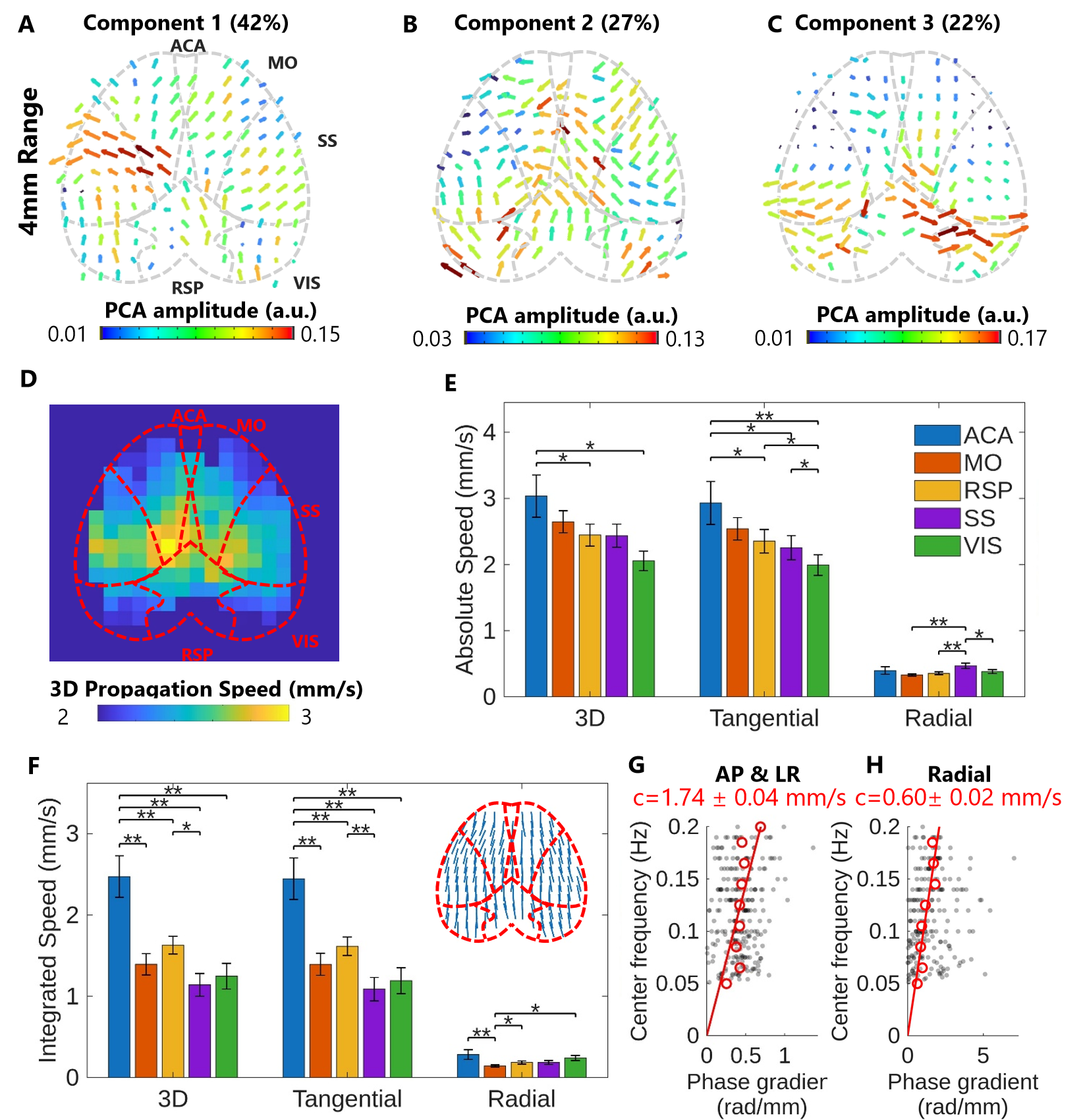

**Supplementary Figure 7. Group-level spatial pattern of 3D vasodynamic propagation speed at large scale.**

(A–C) The top three principal components from PCA of the 3D propagation speed maps at a 4 mm fitting window, explaining 42% (A), 27% (B), and 22% (C) of total variance respectively. Arrow length reflects the tangential projection of the vector magnitude; color encodes PCA amplitude. Cortical region boundaries are shown as dashed lines. (D) Averaged 3D propagation speed amplitude retained at positions passing a one-sample t-test (p < 0.01). (E) Absolute propagation speed for the 3D, tangential, and radial components across five cortical ROIs. At this larger fitting window, ACA exhibited the highest absolute speed in the 3D and tangential components, while SS remained highest in the radial component (repeated-measures ANOVA with Bonferroni post-hoc correction, *p < 0.05, **p < 0.01). (F) Integrated propagation speed for the same three components and ROIs. ACA dominated the integrated 3D and tangential speeds, significantly exceeding all other ROIs (repeated-measures ANOVA with Bonferroni post-hoc correction, *p < 0.05, **p < 0.01). The averaged tangential propagation direction map shows a consistent anterior-to-posterior tendency (Rayleigh test on doubled angle, p < 0.01). Error bars in (E) and (F) represent standard error across animals (n = 18). (G) Scatter plot shows tangential (AP and LR) phase gradients from all 118 trials of 8Hz cortical CBV fMRI in a 1.8x1.8 mm SS ROI, as a function of center frequency, with the estimated propagation speed c = 1.74 ± 0.04 mm/s, R2=0.61 obtained by linear regression. (H) Scatter plot shows radial phase gradients from all 118 trials of 8Hz cortical CBV fMRI in a 2.2 x 2.2 mm SS ROI, as a function of center frequency, with the estimated propagation speed c = 0.60 ± 0.02 mm/s, R2=0.81 obtained by linear regression.

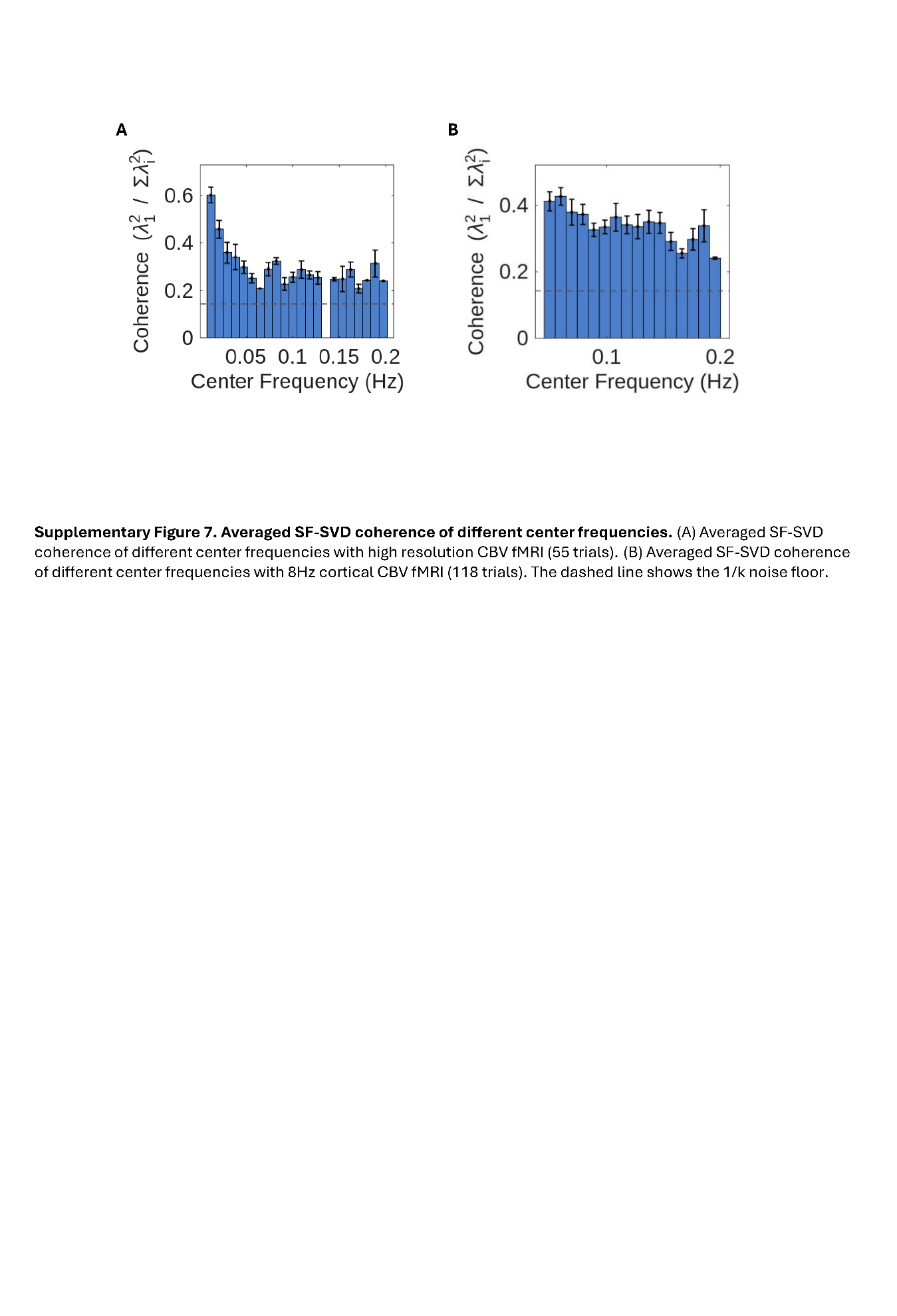

**Supplementary Figure 8. Averaged SF-SVD coherence of different center frequencies.**

(A) Averaged SF-SVD coherence of different center frequencies with high resolution CBV fMRI (55 trials). (B) Averaged SF-SVD coherence of different center frequencies with 8Hz cortical CBV fMRI (118 trials). The dashed line shows the 1/k noise floor.
